# *Candida auris* skin colonization is mediated by Als4112 and interactions with host extracellular matrix proteins

**DOI:** 10.1101/2025.02.13.637978

**Authors:** Guolei Zhao, Jingwen Lyu, Natalia A Veniaminova, Robert Zarnowski, Eliciane Mattos, Chad J Johnson, Derek Quintanilla, Haley Hautau, LeeAnn A. Hold, Bin Xu, Juliet A. E. Anku, Steph S. Steltzer, Kaustav Dasgupta, Darian J. Santana, Ashraf Ibrahim, David Andes, Jeniel E. Nett, Shakti Singh, Adam C. Abraham, Megan L. Killian, J. Michelle Kahlenberg, Sunny Y. Wong, Teresa R. O’Meara

## Abstract

*Candida auris* is an often multidrug-resistant fungal pathogen notorious for persistent skin colonization and transmission in healthcare settings. However, the mechanisms driving its adherence to skin remain poorly understood. Here, we developed *in vitro* systems to allow for detailed analysis of early skin colonization events and identified critical host and pathogen mediators of attachment. Across multiple strains and clades of *C. auris,* we identified that Als4112, a conserved adhesin, is required for skin colonization via keratinocyte attachment and direct interactions with host extracellular matrix (ECM) proteins, especially basement membrane proteins such as laminin. In a murine epicutaneous infection and human skin explants, deletion of *ALS4112* significantly reduced skin colonization, underscoring its essential role in establishing cutaneous persistence. Als4112 also contributes to systemic infection, highlighting the connection between adherence and pathogenicity in this organism. Finally, coating plastic and catheter surfaces with collagen I or III markedly inhibited *C. auris* attachment and biofilm formation, offering an approach to curb nosocomial transmission. Our study highlights the critical role of Als4112 in *C. auris* colonization and virulence *in vivo*, making it an attractive target for future vaccine development. This study also explores the potential of specific collagen coatings as a novel strategy to prevent *C. auris* adherence to abiotic surfaces, offering new therapeutic avenues to control the spread of *C. auris* in healthcare settings.

## Introduction

Since its discovery in 2009 from a Japanese patient ^1^, *C. auris* has rapidly become a significant global health threat. Over the past decade, it has spread to more than 50 countries across six different continents, marking its global emergence as a highly transmissible and multidrug-resistant organism ^2,3^. *C. auris* is predominantly associated with nosocomial outbreaks, raising urgent concerns for public health authorities worldwide ^4–8^. These outbreaks are marked by the pathogen’s ability to persistently colonize patient skin and adhere to abiotic surfaces, creating reservoirs that enable prolonged transmission within healthcare settings ^9–15^.

A key factor contributing to the high transmissibility of *C. auris* is its ability to readily establish high level colonization on the skin and persistently present for months ^5,10,16,17^. Extensive efforts have been made to develop *in vitro* and *in vivo* models to explore the mechanisms of *C. auris* skin colonization. In an *in vitro* model using an artificial human axillary sweat medium, *C. auris* achieves a fungal burden 10 times greater than *Candida albicans* and persists during colonization for over 14 days, which was not observed in *C. albicans* ^18^. This suggests an enhanced ability of *C. auris* to colonize high-sweat body sites, such as the axillae or inguinal crease, which is corroborated by clinical sampling ^10^. While clinical isolates of all four major clades of *C. auris* can colonize the skin ^19^, an *in vivo* murine skin colonization model sampling *C. auris* isolates from different clades demonstrated strain-specific differences in colonization capacity, with one Clade III isolate exhibiting the highest fungal burden 14 days post-inoculation ^20^, Histopathology revealed that *C. auris* can reside deep into the skin, particularly within hair follicles. The fungus persisted in the tissue for up to four months, even when surface swabs yielded negative cultures, suggesting that prolonged colonization is also linked to its ability to survive in deep tissue. These findings likely reflect an advantage for *C. auris* in human skin colonization, and this increased skin colonization has been proposed as a reservoir for nosocomial transmission between patients.

In fungal pathogens, adhesin proteins displayed on the cell surface play crucial roles in surface colonization. In the model fungal pathogen *C. albicans*, targeting adhesin proteins with vaccines results in a decrease in adhesion to and biofilm formation on plastic surfaces, including catheters, invasion of vaginal epithelial cells, and virulence ^21–23^. Crucially, these adhesins mediate interactions with specific host receptors, which facilitates colonization and invasion. For instance, the Als3 adhesin of *C. albicans* binds to receptors on epithelial cells, such as E-cadherin, EGFR, Her2, and c-Met, and receptors on macrophages, such as CR3 ^24–27^. This interplay between fungal adhesins and host receptors highlights key mechanisms of pathogen and host interaction during infection.

Recent research highlights the key role of adhesins in *C. auris* skin colonization and *in vivo* virulence. Our recent study demonstrated the role of two adhesins, Scf1 and Iff4109, in promoting skin colonization and disease ^28^. Additionally, three ALS adhesin proteins were identified in *C. auris* with structural homologies to *C. albicans* Als3. Anti-ALS3 protein antibodies, generated in response to the vaccine containing the N-terminal region of *C. albicans* ALS3 protein, specifically target ALS adhesin proteins on the *C. auris* surface, resulting in disruption of adhesion, biofilm formation, and opsonophagocytic killing ^29^. NDV-3A vaccination significantly improves the protection of mice against hematogenous disseminated *C. auris* infections ^29^. Moreover, Bing and colleagues showed that copy number variation of *ALS4112* accounts for variation in biofilm formation, surface colonization, and virulence among isolates, linking Als4112 amplification to improved skin colonization during natural evolution ^30^. These studies highlight the critical role of adhesins in *C. auris* skin colonization and *in vivo* virulence. However, the specific host cells or proteins that *C. auris* interacts with during skin colonization remain largely unknown, and the exact *C. auris* factors responsible for binding to host components are yet to be identified, underscoring a critical gap in understanding the molecular basis of *C. auris* skin colonization.

To investigate the mechanisms underlying *C. auris* skin colonization, we developed *in vitro* assays to model the adherence between fungi and either keratinocytes or ECM proteins ^31–34^. We identified Als4112 as the primary adhesin mediating keratinocyte adherence across clades, and the expression of this protein was necessary and sufficient for keratinocyte adherence in *C. auris* clinical isolates. Additionally, Als4112 plays a key role in *C. auris* adherence to multiple ECM proteins, such as laminin. Importantly, our *in vitro* findings were corroborated using *in vivo* models, where the deletion of *ALS4112* led to a significant reduction in fungal burden on the skin and virulence during systemic infection. We also discovered that *C. auris* does not adhere to collagen I or III and leveraged this lack of binding to prevent biofilm formation on abiotic surfaces, including on catheters. Overall, our findings suggest Als4112 as a target for vaccine development and collagen coatings as a potential strategy to prevent *C. auris* colonization on abiotic surfaces.

## Results

### *C. auris* adheres to human keratinocyte cells

Keratinocytes are the major epithelial cell type in the skin. Therefore, we established an adherence model using the human immortalized keratinocyte cell line (N/TERT) ^35^ to investigate the ability of *C. auris* to adhere to these cells and assess its requirements for skin colonization. In this model, keratinocytes are incubated with *C. auris* for one hour to allow for adherence, followed by repeated PBS washing to remove non-adherent yeast, fixation, and staining with a FITC-conjugated anti-Candida antibody to identify *C. auris* and CellMask to identify keratinocyte cell bodies. Using this approach, we tested the adherence capacity of representative *C. auris* isolates from four clades (Fig 1A). Consistent with previous reports on plastic adhesion ^28^ and skin adherence ^20^, we observed variation in keratinocyte adherence between these four representative strains (Fig 1A). We then measured the adhesion of 33 *C. auris* isolates representing all five clades, including both clinical isolates and surveillance screening isolates. Importantly, these strains exhibited substantial adhesive variation both within and between clades (*P* < 0.0001, *F* = 8.897) (Fig. 1B). Significant variation in adhesion was observed even between genetically similar isolates of *C. auris*, including the four Clade IV isolates from a single outbreak in Chicago (*P* = 0.0003, *F* = 22.77) (Fig. 1B). This variation potentially indicates variable selection on *C. auris* during an outbreak.

**Fig. 1.**
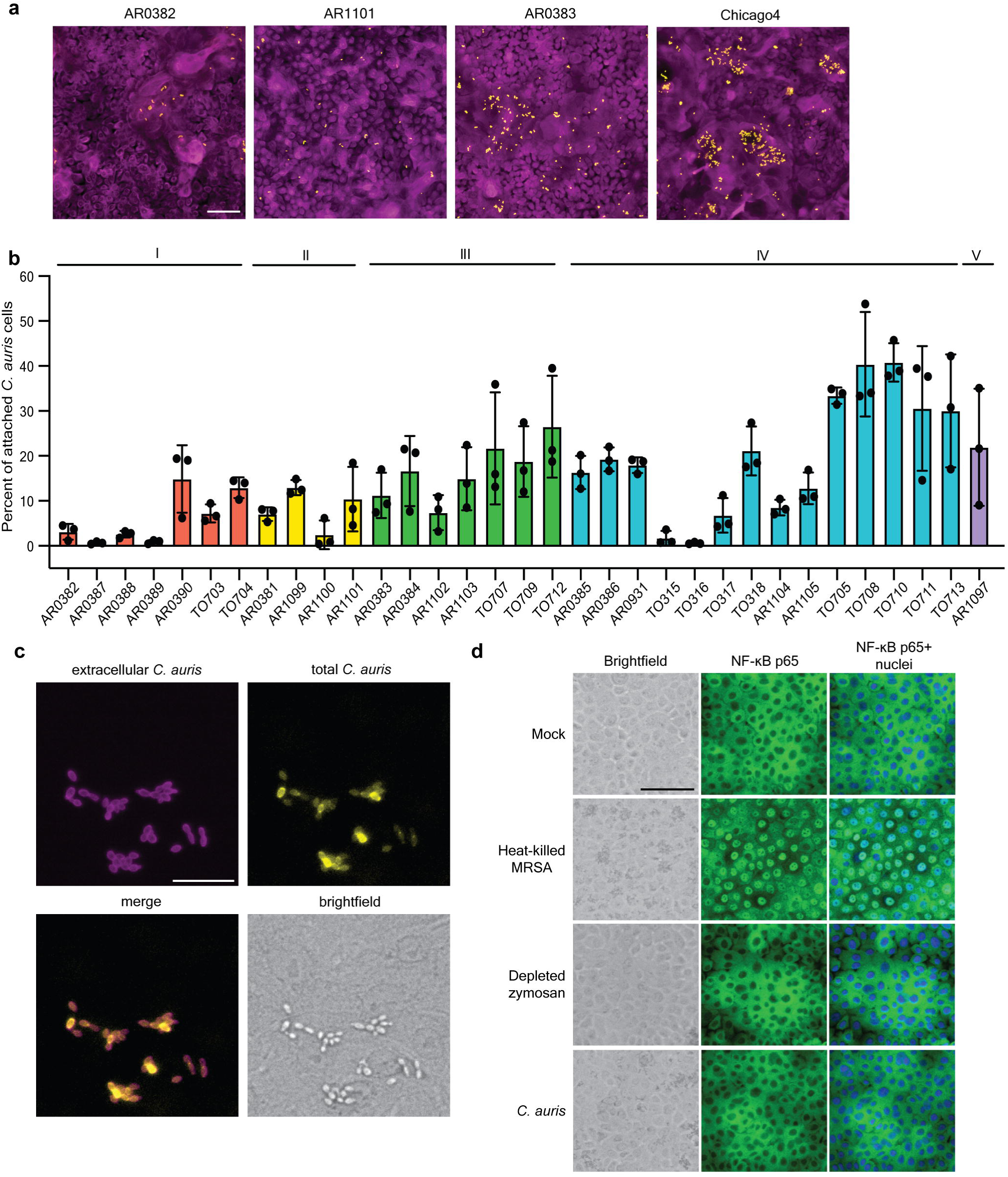
*C. auris* adheres to human keratinocyte cells. **(A)** Representative immunofluorescence images showing adherence of AR0382 (Clade I), AR1102 (Clade II), AR0383 (Clade III), and Chicago4 (Clade IV) cell to human keratinocytes. Magenta = keratinocytes, yellow = *C. auris*. Scale bar, 100 μm. **(B)** Adherence of 33 *C. auris* clinical isolates from five clades to keratinocytes, measured by the proportion of cells remaining attached after washing. Values represent the mean ± s. d., calculated from three biological replicates. **(C)** *C. auris* cells are not internalized. Chicago4 cells adhered to the surface of keratinocytes were detected using an FITC-conjugated anti-*Candida* antibody. Total *C. auris* cells were detected with calcofluor white stain following cell membrane permeabilization. Images are presented in pseudo color. Scale bar, 50 μm. **(D)** *C. auris* does not induce inflammatory signaling. Keratinocytes were unstimulated or stimulated with heat-killed *S. aureus* strain USA300, depleted zymosan, or *C. auris* Chicago4 cells for six hours and nuclear translocation of the NF-LB subunit p65 was detected by fluorescence microscopy. The nuclei were stained with Hoechst (blue). The middle panel shows the NF-kB staining (green) and the right panel shows the overlay. Images are presented in pseudo color. Scale bar, 100 μm. Statistical differences were assessed using one-way ANOVA (B).

It is possible that adherence model would conflate adherence with internalization. However, we differentially stained internal and external fungal cells and observed that the adhering *C. auris* cells remained on the surface of keratinocytes even at four hours of coincubation (Fig. 1C). It is also possible that the adherence is mediated by an active innate immune response from the keratinocytes, which would allow for the expression of molecules that increase binding of the fungal cells. Keratinocytes express Toll-like receptors (TLRs) and C-type lectin receptors (CLRs) to detect and respond to microbial components and drive innate immune responses through the activation of the transcription factor nuclear factor-kappaB (NF-κB) ^36–40^. To determine whether *C. auris* adherence activates immune responses in keratinocytes through NF-κB, we tested the translocation of NF-κB protein from the cytoplasm to the nucleus upon *C. auris* adherence. Heat-killed *Staphylococcus aureus* strain USA300 induced robust nuclear translocation of NF-κB in keratinocytes (Fig. 1D and fig. S1). However, no NF-κB translocation was observed upon treatment with depleted zymosan or *C. auris* infection, even after six hours of coincubation (Fig. 1D and fig. S1). Our results demonstrate that *C. auris* adheres to epithelial keratinocytes without activating strong immune responses and that the binding does not depend on host recognition.

### Als4112 is the major adhesin mediating keratinocyte adherence

We next wanted to identify *C. auris* genes that contribute to keratinocyte adherence using a forward genetic screening approach. To do this, we generated 1,595 insertional mutants in the high-adherent Chicago4 Clade IV strain background using our previously published *Agrobacterium*-mediated insertional mutagenesis approach ^41^. We screened each mutant in this collection in single replicate for those with a significantly decreased level of adherence to keratinocytes as calculated by being 1.8 standard deviations from the plate mean. This revealed three candidates with significant defects (Fig. 2A). We validated these candidates using individual keratinocyte adhesion assays and found that two of them consistently exhibited significant defects in adherence (Fig. 2B). We used whole-genome sequencing and insertion mapping to identify an insertion in the gene *B9J08_004112*, which encodes an ALS adhesin protein on the cell surface, referred to as Als4112 ^28^, and in the gene *B9J08_003899*, which encodes a protein homologous to the *Saccharomyces cerevisiae* Mub1 protein, a subunit of a ubiquitin ligase complex ^42^.

**Fig. 2.**
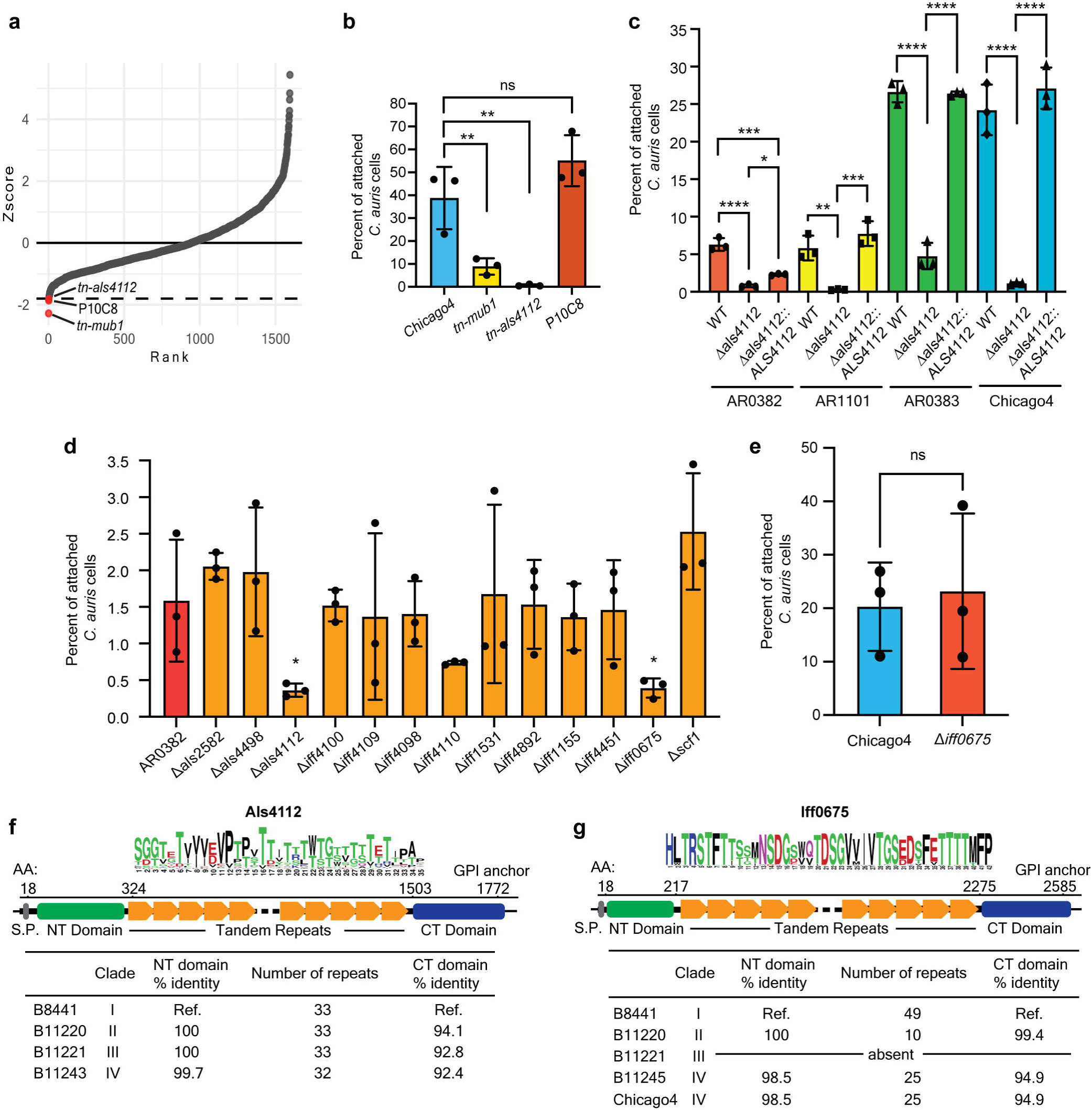
Als4112 is the major adhesin mediating keratinocyte adherence. **(A)** The 1595 insertional mutants in the Chicago4 strain background were screened for adhesion defects by measuring the proportion of cells remaining attached to keratinocytes after three PBS washes. Each dot represents one mutant. Mutants with a Z-score below −1.8 were considered to have a significant adhesion defect. **(B)** Validation of keratinocyte adherence for the three insertional mutants identified in (A). (**C**) Keratinocyte adherence of representative wild-type strains from four clades, Δ*als4112*, and complement strains generated in the same background. **(D)** Keratinocyte adherence of Clade I wild-type AR0382 and mutants lacking one of 13 adhesin-encoding genes in the AR0382 background. (**E**) Adherence of Chicago4 and Δ*iff0675* to keratinocytes was measured by the proportion of cells remaining attached after washing. **(F)** Domain architecture of Als4112 based on the Clade I primary sequence. **(G)** Domain architecture of Iff0675 based on the Clade I primary sequence. Values represent the mean ± s. d., calculated from three biological replicates (B, C, D, and E). Statistical differences were assessed using one-way ANOVA with Dunnett’s multiple comparisons test (B and D), two-way ANOVA with Dunnett’s multiple comparisons test (C), or unpaired t test (E), \**P* ≤ 0.05; \*\**P* ≤ 0.01; \*\*\**P* ≤ 0.001; \*\*\*\**P* ≤ 0.0001; ns: *P* > 0.05.

Given that the *tn-als4112* strain exhibited the most significant decrease in keratinocyte adherence in our screen, and that strains vary in their adhesive capacity, we asked whether the role of Als4112 was conserved across clades. We generated *ALS4112* deletion and complement strains in representative strains from Clades I-IV. The deletion of *ALS4112* led to minimal keratinocyte adherence across all four clades, and the complement strains restored adherence to wild-type levels (Fig. 2C). We then tested whether other adhesins encoded by *C. auris* also contribute to keratinocyte adherence, or if there was specificity in using *ALS4112* during skin colonization. We examined the adherence capacity of 13 individual adhesin mutants generated in the Clade I AR0382 background ^28^. The Δ*als4112* and Δ*iff0675* mutants showed significant decrease compared to the AR0382 parental strain (Fig. 2D), despite an overall lower adherence to keratinocytes in this strain background compared to the AR0383 and Chicago4 strain backgrounds. We then generated an *iff0675* deletion strain in the high-adherence Chicago4 background; however, this did not result in reduced keratinocyte adherence (Fig. 2E), suggesting clade or strain-specific utilization of different adhesin proteins.

To clarify the distinct roles of Als4112 and Iff0675 in mediating adherence, we analyzed the conservation and the domain architecture of these proteins across clades. Als4112 displayed high conservation in both its N-terminal and C-terminal regions, as well as in the number of tandem repeats (Fig. 2F). In contrast, while Iff0675 also showed strong conservation in its N- and C-terminal domains, the number of tandem repeats varied significantly among clades (Fig. 2G), and the Clade III strain B11221 (AR0383) lacked Iff0675, but showed high adherence to keratinocytes (Fig. 2C). Moreover, while the Iff0675 protein has more conservation in its repeat sequence than Als4112, there was more variance in the number of repeats between clades. Together, these results suggest that Als4112 is the major adhesin required to mediate keratinocyte adherence across clades, although there is variability between strains on which adhesins are used for binding. Therefore, we focused on the role of Als4112 in skin colonization of *C. auris*.

### Modulating Als4112 expression levels regulates *C. auris* adherence to keratinocytes

To investigate the general reliance on Als4112 and start to explain the variability among *C. auris* strains in keratinocyte adherence, we compared the transcript abundance of *ALS4112* with the keratinocyte adherence of our 33 clinical isolates. We observed a strong positive correlation between *ALS4112* transcription abundance and adhesion across isolates, regardless of clade (*r*=0.7441, *P*<0.0001) (Fig. 3A). This indicates that adhesive capacity among *C. auris* isolates is associated with *ALS4112* expression.

**Fig. 3.**
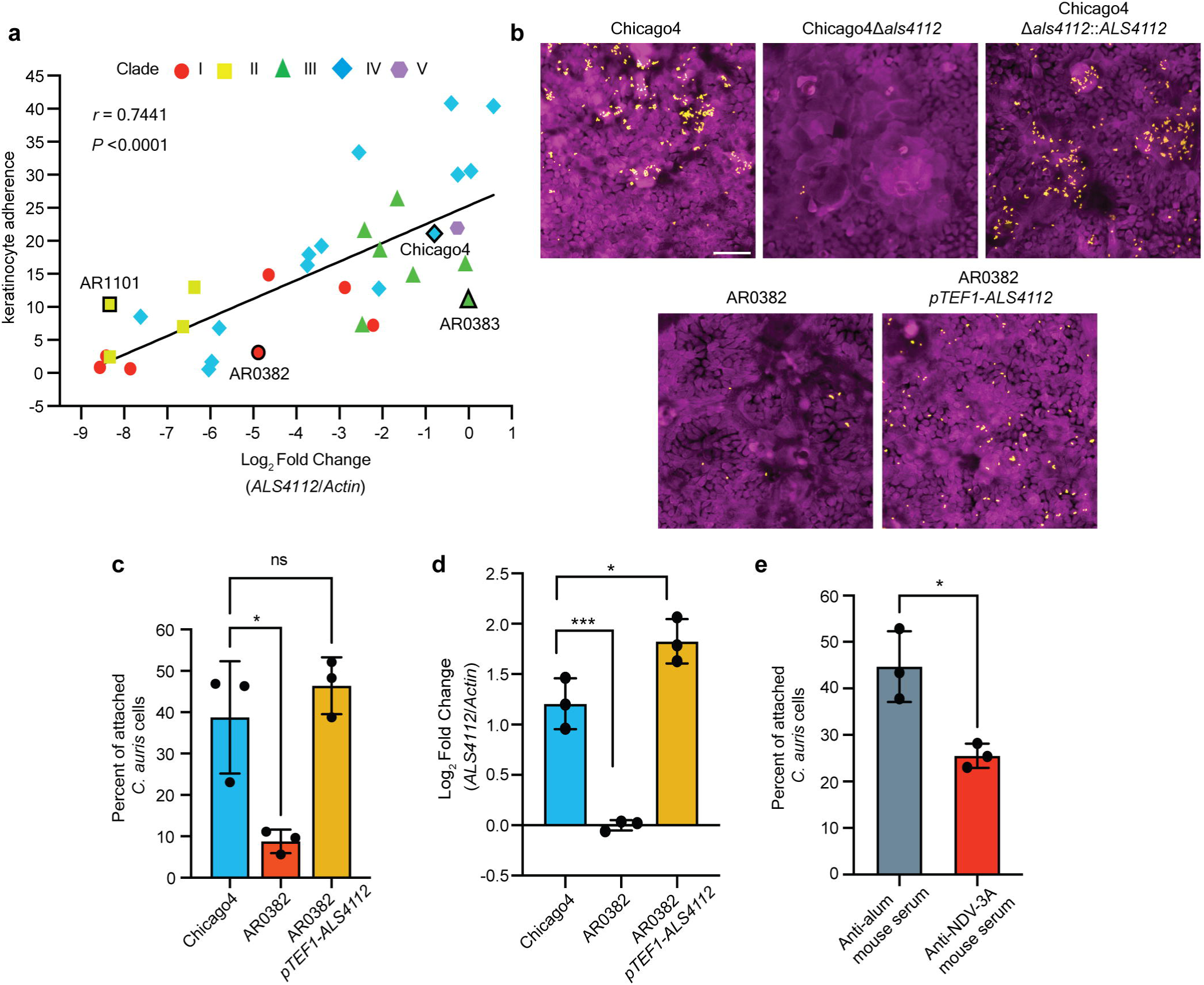
Modulating Als4112 expression levels regulates *C. auris* adherence to keratinocytes. **(A)** *ALS4112* transcript abundance is associated with adhesion to keratinocytes in the same 33 *C. auris* isolates from (1B). Log_2_ fold change values are expressed relative to AR0383 and normalized to *Actin*. Each point signifies the mean of three biological replicates. Pearson correlation coefficient and *P* value are indicated. **(B)** Representative immunofluorescence images showing keratinocyte adherence of Chicago4, Δ*als4112*, and complement strains generated in the Chicago4 background, as well as AR0382 and AR0382 *pTEF1*-*ALS4112* strains. Magenta = keratinocytes, yellow = *C. auris.* Images are presented in pseudo color. Scale bar, 100 μm. **(C)** Overexpression of *ALS4112* using *pTEF1* promoter significantly enhanced keratinocyte adhesion in the poorly adhesive AR0382 background. **(D)** *pTEF1* promoter significantly increased the transcript level of *ALS4112* in AR0382 background, as measured by qRT-PCR. **(E)** Chicago4 cells pretreated with anti-NDV-3A serum exhibited reduced adherence to keratinocytes compared to cells pretreated with control serum. Values represent the mean ± s. d., calculated from three biological replicates (C, D, and E). Statistical differences were assessed using one-way ANOVA with Dunnett’s multiple comparisons test (C and D) or unpaired t test (E); \**P* ≤ 0.05; \*\*\**P* ≤ 0.001; ns: *P* > 0.05.

To test whether overexpression of *ALS4112* is sufficient to enhance keratinocyte adherence, we overexpressed *ALS4112* in the otherwise low-adherence AR0382 background using the *pTEF1* strong constitutive promoter. Transcriptional overexpression of *ALS4112* was sufficient to elevate adhesion of the low adherence AR0382 to levels similar to that of the Chicago4 strain (Fig. 3B and 3C). The *ALS4112* transcription level driven by the *pTEF1* promoter was also comparable to that of the Chicago4 strain (Fig. 3D). This suggests that Als4112 is sufficient to allow for keratinocyte adherence.

To further investigate the role of ALS adhesins in keratinocyte adherence and explore potential therapeutic interventions, we examined the effects of antibodies targeting ALS proteins. The NDV-3A vaccine is based on the N-terminus of *C. albicans* Als3 protein, and IgG from NDV-3A immunized mice binds to *C. auris* cell surfaces, likely targeting *C. auris* ALS adhesins ^29^. We treated Chicago4 cells with serum from NDV-3A- or alum-immunized mice and tested keratinocyte adherence of the pretreated cells. We observed a ∼30% decrease in keratinocyte adherence when Chicago4 cells were pretreated with serum from NDV-3A-immunized mice compared to those treated with serum from alum-immunized mice (Fig. 3E). These findings strongly suggest that antibodies targeting ALS adhesins can significantly inhibit keratinocyte adherence of *C. auris*.

### Als4112 mediates *C. auris* adhering to ECM proteins

*C. auris* can reside on the skin and within murine hair follicles ^20^. Human hair follicles are embedded in the dermis, which contains a complex mixture of ECM proteins, including laminin ^34^, which are essential for the structural integrity and function of the skin. We, therefore, wanted to investigate whether *C. auris* binds directly to ECM proteins to mediate colonization. To test this hypothesis, we leveraged microscopy approaches to directly measure *C. auris* attachment to a glass slide printed with 35 distinct ECM protein combinations on a hydrogel surface. We observed that *C. auris* exhibited strong binding to tropoelastin, collagen V, and laminin, but not significant adherence to collagen I, III, IV, VI, fibronectin, or vitronectin (Fig. 4A and 4B). This indicated specificity in the attachment patterns of *C. auris* to different ECM proteins, including specific structural variants of collagens.

**Fig. 4.**
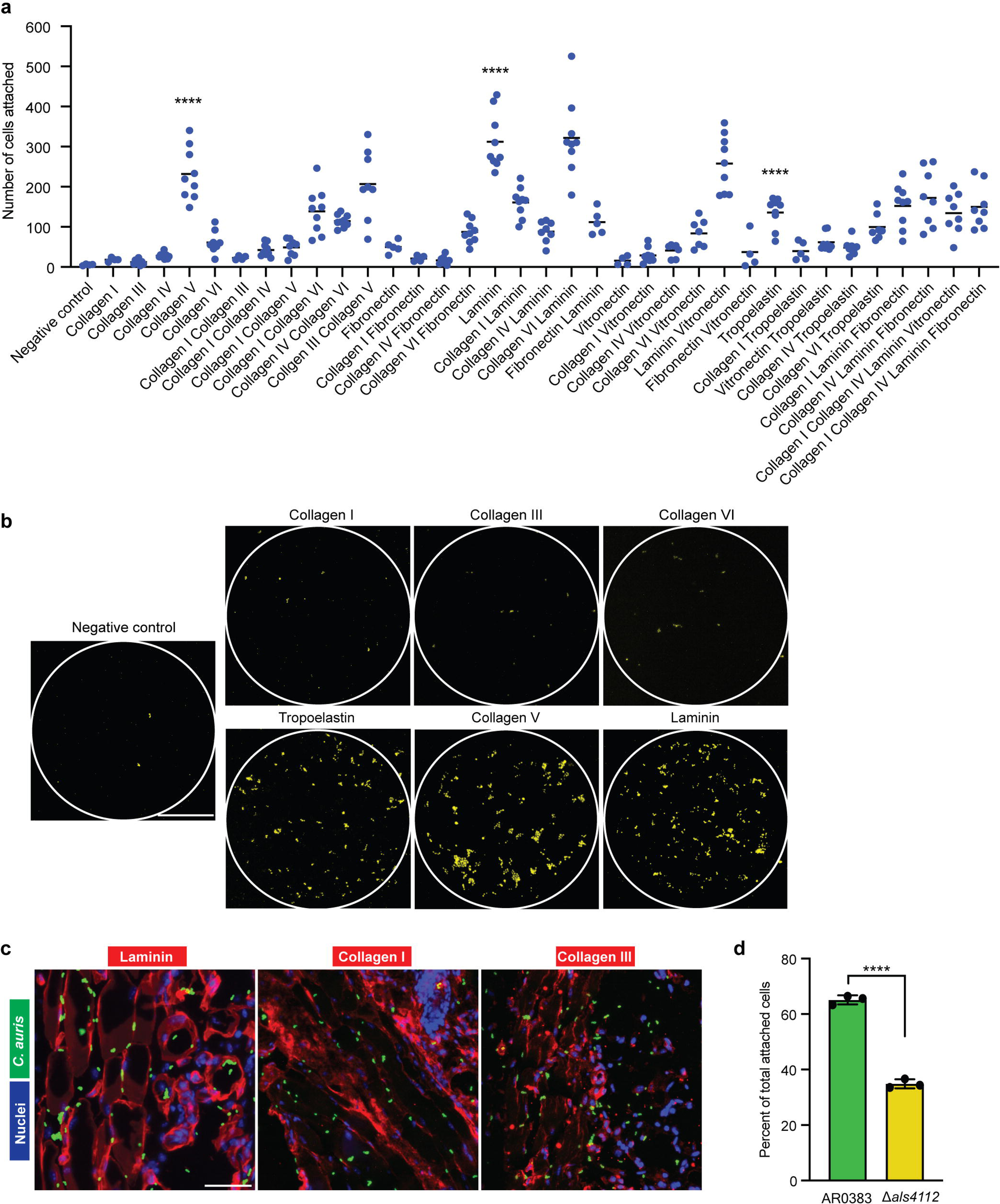
Als4112 mediates *C. auris* adherence to ECM proteins. **(A)** The number of cells attached to human ECM protein prints, counted with CellProfiler. **(B)** Fluorescence microscopy images showing adhesion of AR0383 Eno1-RFP cells to human ECM protein prints, highlighted by circles. Yellow dots represent *C. auris* cells, shown in pseudo color. Scale bar, 100 μm. **(C)** In situ incubation of *C. auris* on mouse skin shows co-localization with laminin, but not collagen I or III. Scale bar, 50 μm. **(D)** The number of AR0383 Eno1-RFP and AR0383 Δ*als4112* cells that remained attached to laminin-coated glass coverslips after washing were counted. The percentage of attached cells for each strain relative to the total number of attached cells was plotted. Values represent the mean ± s. d., calculated from three biological replicates. Statistical differences were assessed using one-way ANOVA with Dunnett’s multiple comparisons test (A) or unpaired t test (D); \*\*\*\**P* ≤ 0.0001.

To determine whether *C. auris* exhibits direct adhesion to ECM proteins within the native skin environment, we performed an *in situ* binding assay on murine skin slices. Our analysis revealed that *C. auris* exhibited strong colocalization with skin laminin but not with collagen I or III (Fig. 4C), suggesting a specific binding affinity for laminin in the skin. Given that laminin is a principal ECM component in the skin ^43^, we performed a competition adhesion assay to evaluate whether Als4112 plays a role in mediating *C. auris* binding to laminin. Fluorescently labeled AR0383 cells were mixed in a 1:1 ratio with Δ*als4112* cells in the same background and incubated on laminin-coated coverslips. The number of cells remaining attached to the surfaces after washing was counted to assess binding efficiency to laminin. The Δ*als4112* strain demonstrated a 50% reduction in adherence to laminin-coated coverslips (Fig. 4D), underscoring the crucial role of Als4112 in facilitating ECM adherence in *C. auris*. Our findings indicate that *C. auris* binds to specific skin ECM proteins and highlight Als4112 as a critical factor facilitating the pathogen’s interaction with key ECM components in skin tissue.

### Als4112 facilitates *C. auris* colonization of host skin

Given the robust skin colonization ability of *C. auris*, we utilized a murine epicutaneous skin colonization model to examine its distribution on the skin and identify colonization patterns that may contribute to its transmissibility and persistence. C57BL/6 mice were inoculated with the high-adherent Chicago4 strain on depilated dorsal skin. Skin tissue was collected either two hours after infection to evaluate early colonization or two days after infection to examine established colonization. Using immunofluorescent staining, we discovered that Chicago4 cells rapidly established themselves on the murine skin surface within just two hours (Fig. 5A). By two days post-infection, *C. auris* had formed a dense coating on the skin surface, robustly coating the hair shafts and residing in the hair follicle (Fig. 5A). Strikingly, the infection had progressed further, with Chicago4 cells invading the skin tissue, infiltrating both the epidermis and penetrating deep into the dermis (Fig. 5A). To further assess potential tissue damage or inflammation caused by *C. auris* infection, we performed Hematoxylin and Eosin (H&E) staining on infected skin tissues. Histopathological analysis revealed no apparent lesions or significant structural disruptions in the infected skin (Fig. 5B). Consistent with our immunofluorescent staining results, we observed robust *C. auris* colonization on the skin surface, also with fungal cells densely coating the hair shaft (Fig. 5B). These findings indicate that while *C. auris* establishes extensive skin colonization, it does not appear to cause overt inflammatory damage.

**Fig. 5.**
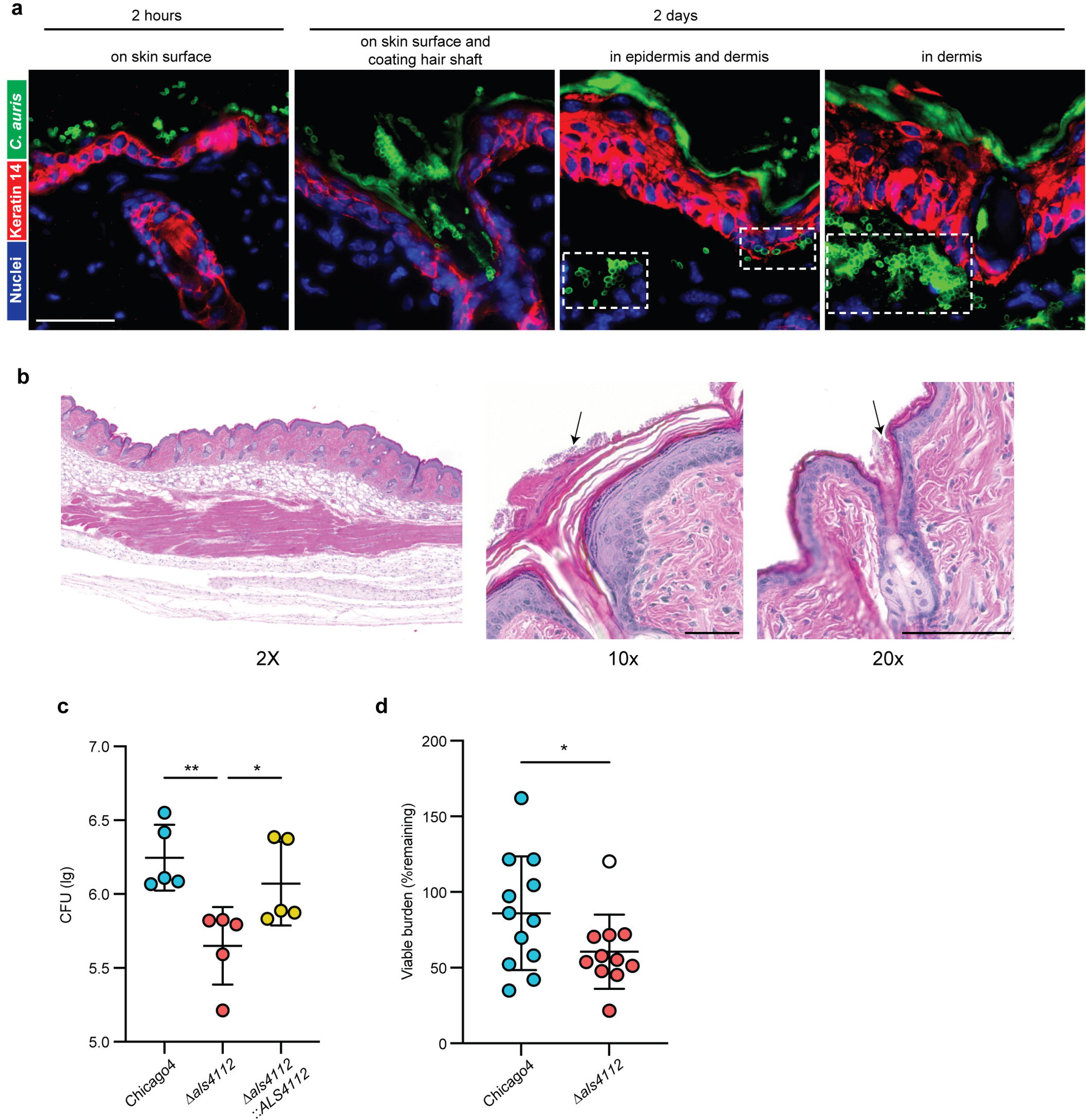
Als4112 facilitates *C. auris* colonization of host skin. **(A)** Immunofluorescence staining was performed to visualize the localization of *C. auris* cells (green) in the skin 2 hours or 2 days post infection. The keratin 14 signal (red) marked the epidermis and Hoechst was used to stain the nuclei. Scale bar, 25 μm. **(B)** H&E staining of murine skin infected with *C. auris*. Arrows point to fungal cells on the skin surface and coating the hair shaft. Scale bar, 100 μm **(C)** *C. auris* bioburden after 2 days of incubation on murine skin *in vivo*. Data are presented as CFU/g of tissue, normalized to the delivered inocula. n = 5 mice. **(D)** *C. auris* cells were applied to *ex vivo* human skin samples and incubated for 24 hours before washing. Recovered CFU after wash were normalized to the average total burden for each strain in each experiment (% remaining). Values represent the mean ± s. d., calculated from two biological replicates with 5-6 technical replicates each, depending on skin availability. A single outlier (open circle) was identified and removed using Grubb’s test. Excluding the outlier point, *P* = 0.02, including the outlier point, *P* = 0.07. Statistical differences were determined using one-way ANOVA with Dunnett’s multiple comparisons test (C) or unpaired t-test (D); \**P* ≤ 0.05; \*\**P* ≤ 0.01.

Thus far, our data demonstrated that Als4112 is critical for binding to *in vitro* proxies for colonization. We then used our murine skin colonization model to investigate whether Als4112 is essential for skin colonization *in vivo*. The Chicago4 wild-type strain, the Δ*als4112* mutant, or the complement strain were inoculated on depilated dorsal skin of C57BL/6 mice. After 2 days, skin tissue was harvested, homogenized, and fungal burden was quantified by colony-forming units (CFU). The wild-type Chicago4 strain exhibited a high fungal burden, with an average log_10_ CFU of around 6.3 (Fig. 5C). Strikingly, the Δ*als4112* mutant showed a dramatic and significant reduction in fungal burden, with an average 7-fold decrease in colonization capacity (Fig. 5C). Importantly, the complementation strain restored fungal burden to levels comparable to the wild-type strain. A similar pattern was observed in the *ex vivo* human skin colonization model, where the Δ*als4112* mutant demonstrated a diminished ability to colonize human skin compared to the Chicago4 strain (Fig. 5D). The reduction in fungal burden observed in the Δ*als4112* mutant was not due to a growth defect, as the mutant displayed growth comparable to that of the wild-type and complement strains (fig. S2A). Our data underscore Als4112 as a crucial factor for efficient *C. auris* skin colonization.

### Als4112 promotes *C. auris* virulence during systemic infection

Given the potential for Als4112 to interact with host cells and tissues, we further explored its role in systemic infections. Immunosuppressed CD-1 mice were infected with a lethal inoculum of the Chicago4 wild-type strain, the Δ*als4112* mutant, or the complemented strain and monitored for survival and histopathological changes in primary target organs (kidney and heart). Mice infected with the wild-type strain exhibited significant weight loss by day 7, whereas mice infected with the Δ*als4112* mutant did not (Fig. 6A). Furthermore, the Δ*als4112* mutant demonstrated significantly reduced virulence, with 60% survival and a median survival time exceeding 21 days, compared to the wild-type strain, which caused 100% mortality and a median survival time of 11.5 days (*P* = 0.0008) (Fig. 6A). The complemented strain partially regained virulence, resulting in reduced survival compared to the Δ*als4112* mutant (40% vs. 60%, *P* = 0.2754) (Fig. 6A). Kidney and heart tissues were collected seven days after intravenous *C. auris* infection for histopathological analysis by H&E staining. Deletion of *als4112* resulted in reduced lesions in these organs (Fig. 6B and 6C). The complement strain exhibited a partial restoration of virulence and a higher number of lesions compared to the *als4112* mutant, indicating a role for Als4112 in facilitating tissue colonization (Fig. 6B and 6C).

**Fig. 6.**
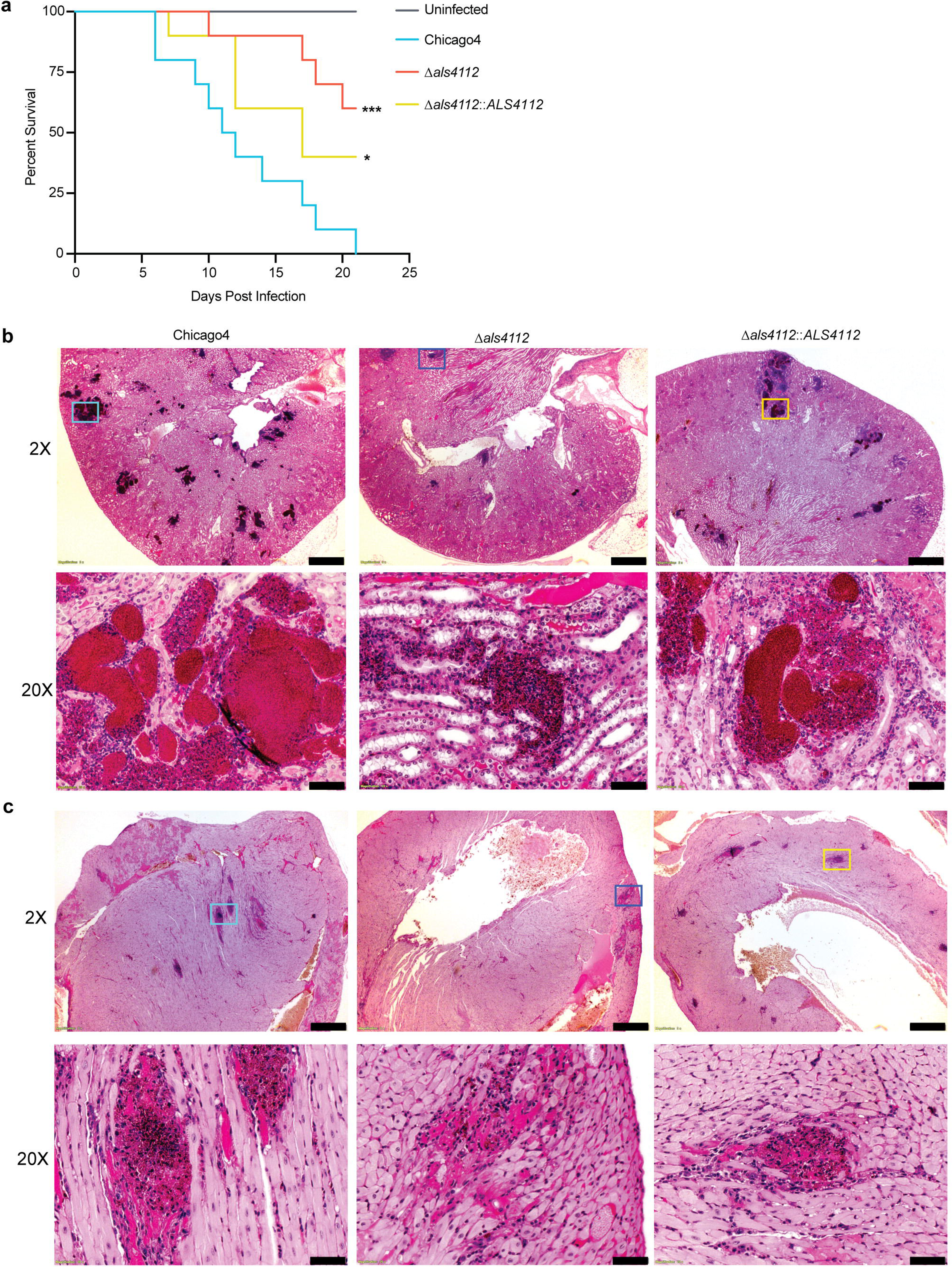
Als4112 promotes *C. auris* virulence during systemic infection. **(A)** Immunosuppressed mice were infected intravenously with 5 × 10^7^ *C. auris* cells. Ten infected mice for each group were monitored for survival for 21 days. **(B)** Kidney and **(C)** Heart of the *C. auris*-infected mice were sectioned and stained with H&E. Stained slides were imaged at 2X and 20X resolution using Olympus bright field microscopy. The lower panel for each organ represents the magnified area of colored boxes from 2X panel. Scale bar, 500 μm (2X) and 50 μm (20X). Statistical differences were determined using Log-rank (Mantel-Cox) test (A); \**P* ≤ 0.05; \*\*\**P* ≤ 0.001.

Due to the incomplete complementation of virulence, we performed a comparative analysis of transcript abundance for *ALS4112* and other adhesin-encoding genes in the *ALS4112* complement strain. Our results showed that although *ALS4112* transcription in the complement strain was comparable to that of the wild-type strain, transcription of *IFF4109* was absent (fig. S2B). This gene has previously been shown to play a role in *C. auris* systemic infection ^28^. Additionally, the gene *IFF4110* was lost (fig. S2B), further highlighting a disruption in adhesin gene expression. *IFF4109*, *IFF4110*, and *ALS4112* are tightly clustered within a 23 kb region near the telomere ends. Specifically, *ALS4112* and *IFF4110* are separated by 4.3 kb, while *IFF4110* and *IFF4109* are only 1.9 kb apart. This dense arrangement near the telomere may make the region unstable during genetic manipulation. These findings suggest that the partial recovery of virulence observed in the complement strain during systemic infection may be attributable to the dysfunction of these other important adhesin-encoding genes. Taken together, our data suggest Als4112 is essential for systemic infection *in vivo*, while also revealing the potential contribution of other adhesins to *in vivo* virulence.

### Coating plastic surfaces with collagen I or III inhibited *C. auris* colonization

*C. auris* is known for its ability to adhere to synthetic surfaces, posing a significant threat to patients with indwelling devices, such as catheters ^11,12,28,44^. Given this context, our ECM adhesion experiment revealed that *C. auris* does not bind to collagen I or III, which prompted us to investigate whether coating plastic surfaces with collagen I or III could inhibit *C. auris* adherence. To test this, we first coated 96-well plates with collagen I or III and measured the adherence of four representative *C. auris* clinical isolates from four clades. While *C. auris* exhibited strong adherence to uncoated plastic surfaces, coating them with collagen I or collagen III effectively inhibited attachment (Fig. 7A). Remarkably, this reduction in adhesion was observed across all four clades of *C. auris*, underscoring the broad efficacy of specific collagen coatings in preventing colonization, although the extent of reduction varied among clades (Fig. 7B). The reduced adherence of *C. auris* on collagen I- or III-coated plastic surfaces appears to be specific, as BSA coating did not inhibit *C. auris* adherence across different clades (fig. S3A). Our observations highlight the potential for collagen coatings to serve as a universal strategy for mitigating *C. auris* adherence, regardless of clade variability.

**Fig. 7.**
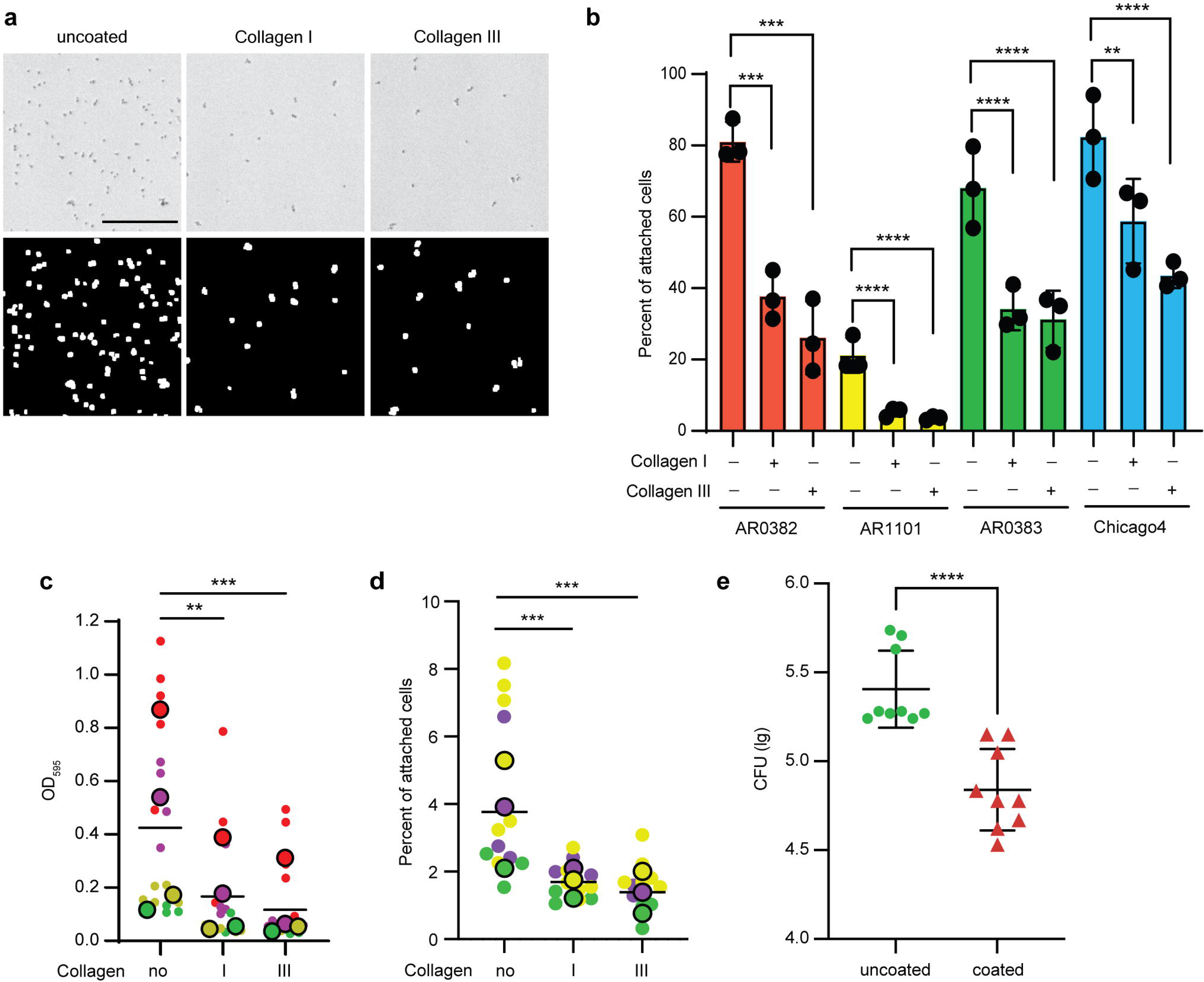
Coating surfaces with collagen I or III inhibits *C. auris* colonization. **(A)** Top: Brightfield images showing AR0383 cells remaining attached to uncoated or collagen-coated 96-well plates after three washes with PBS. Bottom: Images processed by ImageJ for cell counting with CellProfiler. Scale bar, 100 μm. **(B)** Adhesion of AR0382, AR1101, AR0383, and Chicago4 cells to uncoated or specific collagen-coated 96-well plates, measured by the proportion of cells remaining attached after washing. **(C)** Biofilms were grown on uncoated or specific collagen-coated 96-well plates in RPMI at 37 °C for 24 hours. Quantification of biofilm formation was determined with crystal violet staining. (**D**) Adhesion of AR0383 to catheter was measured by the proportion of cells remaining attached after washing. Superplots showing the summary of data from biological replicates, incorporating all associated technical replicates for statistical analysis (C and D). (**E**) Uncoated or collagen III-coated polyethylene central venous catheters were inserted into the jugular veins of rats and inoculated intraluminally with *C. auris* AR0383 strain. CFU was used to assess *C. auris* cells colonized on the luminal catheter surface 24 hours after infection. Plot showing the summary of data from two independent experiments, incorporating all associated technical replicates for statistical analysis, n = 3 rats. Statistical differences were determined using one-way ANOVA with Dunnett’s multiple comparisons test (B, C, and D) or unpaired t test (E); \*\**P* ≤ 0.01; \*\*\**P* ≤ 0.001; \*\*\*\**P* ≤ 0.0001.

To further explore the impact of collagen coating, we assessed its effect not only on the initial adhesion of *C. auris* but also on its subsequent biofilm formation, a crucial factor in the persistence of infections and resistance to treatment ^12,45–48^. We assessed biofilm development of *C. auris* Clade III strain AR0383, which showed high adherence on plastic surfaces. Using crystal violet and calcofluor white staining, we observed a significant reduction in biofilm formation by *C. auris* AR0383 on collagen I- or III-coated 96-well plates (Fig. 7C and fig. S3B). This reduction in biofilm formation was observed across strains from other clades as well (fig. S3B), suggesting that collagen coatings effectively inhibit biofilm development on plastic surfaces. To apply these findings to clinically relevant materials, we next tested *C. auris* adherence on polyethylene (PE) catheters coated with collagen I or III. Consistent with previous results, the AR0383 strain showed significantly reduced adhesion to collagen-coated catheters (Fig. 7D). This pattern was also observed *in vivo*, where the high-adherent strain AR0383 exhibited an average 6-fold decrease in colonizing the luminal surface of collagen III-coated PE catheters in a rat central venous model compared to uncoated controls (Fig. 7E). These results further underscore the potential of specific collagen coatings in reducing *C. auris* biofilm formation on medical devices.

## Discussion

The highly transmissible fungal pathogen *C. auris* can readily colonize human skin and shedding from patient skin is considered one of the reservoirs for nosocomial contamination by *C. auris* ^10,16^. Our study demonstrates that *C. auris* adheres to both keratinocytes and ECM proteins found in the skin. The ability of *C. auris* to adhere to multiple skin components may explain its capacity for high levels of skin colonization.

We observed strain-specific variation in human keratinocyte adherence among *C. auris* clinical isolates, which aligns with previous findings of the varied skin colonization abilities in a murine model ^20^. However, Eix et al. did not report such variation using a porcine skin colonization model ^19^. This inconsistency may be due to the specific strains tested, or the use of different models, where variations in skin structure could have influenced the results. Despite the strain-specific variation in keratinocyte adherence, we identified Als4112, a surface adhesin, as the primary mediator of keratinocyte adherence across clades. This critical role of Als4112 in skin colonization was further validated in our *in vivo* murine skin colonization model. Our data also revealed a positive correlation between the transcriptional abundance of *ALS4112* and the level of keratinocyte adherence among clinical isolates, suggesting that differences in Als4112 expression may partially explain the strain-specific variation in skin colonization potential. This is consistent with a previous report showing a correlation between genomic amplification of *ALS4112* and enhanced skin colonization in *C. auris* clinical isolates ^30^. These findings further suggest that understanding how *ALS4112* transcription is regulated could provide valuable insights into the mechanisms controlling skin colonization, revealing new ways to disrupt *C. auris* colonization.

We found that the role of Iff0675 in mediating keratinocyte adherence varied in different strains. The comparison of Als4112 and Iff0675 in their function and domain architecture suggests that the number of tandem repeats in adhesin proteins, rather than the repeat sequence, may play a key role in their functions. This agrees with previous studies showing that variations in tandem repeat copy numbers within fungal adhesin proteins can affect their surface hydrophobicity, cell surface display, and protein shedding, ultimately influencing their functions ^49,50^. This suggests that the function or utilization of specific proteins may vary depending on the genetic background. Therefore, it will be crucial to consider potential strain-specific differences when studying the function of proteins.

Previously, we reported that two *C. auris* adhesins, Scf1 and Iff4109, contribute to skin colonization ^28^, but their deletion did not reduce adhesion in our keratinocyte assay—suggesting these adhesins may function independently of direct keratinocyte binding. One limitation of our model is the inability of isolated keratinocytes to represent the complexity of different skin layers. The skin epidermis consists of several keratinocyte layers with varying differentiation states, with the outermost layer composed of anucleate, dead keratinocytes. Our keratinocyte model cannot fully capture the spectrum of differentiated keratinocytes, nor the complete array of ECM proteins present in skin tissue. Nevertheless, the decreased keratinocyte adherence observed for the *als4112* mutant aligns with our *in vivo* murine model, where reduced skin colonization was also evident. Thus, the keratinocyte adhesion model remains a useful tool to begin exploring the genetic mechanisms underlying *C. auris* skin colonization.

In addition to human keratinocytes, we observed that Als4112 mediates adherence to specific ECM proteins, specifically those associated with the basement membrane. Recognition of ECM components by adhesins is far from unique. In the bacterial pathogen *Klebsiella pneumoniae*, the MrkD adhesin specifically binds to collagen V, enabling adhesion to tubular basement membranes, Bowman’s capsule, arterial walls, and the interstitial connective tissues of the kidney. ^51–54^. In *Escherichia coli*, various adhesins from different fimbrial types facilitate binding to host ECM proteins, such as collagens and fibronectin, which contributes to its persistent colonization and infection ^55–59^. In the fungal pathogen, *C. albicans*, Als3 facilitates adhesion to fibrinogen and collagen IV ^60^. The critical role of Als4112 in binding both keratinocytes and ECM proteins likely explains its importance in skin colonization in our *in vivo* model. Moreover, our observation that blocking *C. auris* with anti-NDV-3A serum inhibited keratinocyte adherence further supports Als4112 as a promising target for vaccination or therapeutic strategies aimed at reducing *C. auris* skin colonization.

Our finding that *C. auris* exhibits varying affinities for different ECM proteins, particularly the various types of collagens, is intriguing. Among the collagens tested, collagen I, III, and V are classified as fibril-forming collagens, while collagen IV forms network structures, and collagen VI forms beaded filaments ^61^. Despite these structural differences, *C. auris* showed a notably high affinity only for collagen V. This observation suggests that the macro-structural classification of collagens may not be the primary factor determining their interaction with *C. auris*. Instead, other factors such as the specific molecular composition or differing surface properties could be driving the adherence. Many fungal pathogens are known to degrade ECM components as part of their tissue invasion strategy ^62–65^. Although *C. auris* does not bind to certain ECM proteins, investigating whether it can degrade ECM proteins could provide valuable insights into its mechanisms of tissue invasion and pathogenesis. Additionally, exploring whether different *C. auris* isolates exhibit preferences for specific ECM proteins could enhance our understanding of strain-specific virulence factors.

We investigated the distribution of *C. auris* during skin colonization and observed its presence across various skin layers. Our findings show that *C. auris* rapidly establishes itself on the murine skin surface after inoculation, demonstrating its ability to readily adhere and initiate colonization. This observation aligns with previous reports of *C. auris* residing on the skin surface and within hair follicles ^20,66^. As colonization progresses, we found that *C. auris* not only forms a dense layer on the skin surface but also penetrates deeper into the tissue, infiltrating the epidermis and extending into the dermis, suggesting its capacity for deeper tissue invasion. However, skin invasion was not reported in previous studies ^20,66^, which may be due to differences in the *C. auris* strains used in *in vivo* skin colonization experiments. In this study, we utilized the Chicago4 strain with high keratinocyte adherence, whereas prior reports used Clade I strains, which generally exhibit low adherence to keratinocytes. Notably, strains from different clades have been reported to vary in their skin colonization capacity ^20^. Overall, the presence of *C. auris* on the skin surface and deep in skin tissue may explain its ability to persist and spread in clinical settings, while also increasing the risk of systemic infections. Furthermore, we identified the cell surface adhesin Als4112 as a key factor in the ability of *C. auris* to colonize skin. These findings emphasize the importance of studying the mechanisms underlying *C. auris* colonization and invasion to better understand its role in infections and outbreaks.

The deletion of the Als4112 gene also significantly reduced the virulence of *C. auris* in a systemic candidiasis model, suggesting its role in host tissue invasion. This finding corroborates the previous work by Singh et al., where the NDV-3A vaccine targeting ALS orthologs, including the *C. albicans* Als3p protein, protected mice from systemic infection ^29^. Our results reaffirm the involvement of ALS adhesins in *C. auris* virulence *in vivo* and highlight Als4112 as a potential vaccine target.

A key finding in this study is that collagen I or III coatings can reduce *C. auris* colonization and biofilm formation on plastic surfaces, including catheters, in both *in vitro* and *in vivo* models. This is particularly significant given the persistence of *C. auris* on abiotic surfaces in healthcare settings, where it plays a major role in nosocomial outbreaks. Notably, collagen I may have a dominant effect in blocking *C. auris* adherence to high-adhesion ECM proteins, such as laminin, collagen V, and tropoelastin. We observed a reduced number of *C. auris* cells adhering to these ECM proteins when collagen I was applied. Our findings offer a practical, safe, sustainable, and cost-effective solution to minimize *C. auris* contamination and colonization on medical devices, such as catheters. Further studies are needed to evaluate the broader effects of collagen or other biomaterials in limiting surface colonization by other pathogens.

In conclusion, our study provides significant insights into the mechanisms of *C. auris* skin colonization, identifying Als4112 as a critical adhesin mediating adherence to both keratinocytes and ECM proteins. By demonstrating the role of Als4112 in skin colonization and its potential as a therapeutic target, this work lays the foundation for future strategies aimed at preventing *C. auris* colonization and reducing the risk of nosocomial transmission. Additionally, our findings on the use of collagen coatings to inhibit *C. auris* biofilm formation offer a promising approach to mitigating colonization on medical devices, providing a new avenue for infection control in healthcare settings.

## Materials and Methods

### Strains and Growth Conditions

A list of all strains used in this study is provided in Table S1. Clinical *C. auris* isolates were obtained from the CDC/FDA Antibiotic Resistant Isolate Bank ^67^, were generously provided by Mary Hayden from Rush University Medical Center, Chicago, IL or the University of Michigan Infection Prevention team. All *C. auris* strains were stored in 25% glycerol stocks at −80°C. For adhesion assays, cells grown in liquid YPD (1% yeast extract, 2% peptone, 2% glucose) overnight at 30°C were used.

### Primers and Plasmids

The plasmids used in this study are listed in Table S2, and the primers used are listed in Table S3. Transformation cassettes for *C. auris* were maintained within the multiple cloning site of the pUC19 cloning vector and assembled from fragments using the NEBuilder HIFI DNA Assembly Master Mix (NEB #E2621), according to the manufacturer’s instructions.

### Plasmid and Strain Construction

*C. auris* strains were generated with a transient CRISPR system ^41^. The universal *CAS9* cassette was amplified from plasmid pTO135 using oTO143-oTO41. To make *als4112* mutant strains, the gRNA cassette was constructed using a Splice-On-Extension reaction, incorporating a 20-bp specific gRNA targeting the *ALS4112* gene in primers oTO1486-oTO1487. The repair template was amplified from plasmid pTO166 using oTO18-oTO19.

To make plasmid to add back *ALS4112*, the plasmid backbone was amplified from pTO139 using oTO590-oTO591. The ORF for *ALS4112* (*B9J08_004112*) along with approximately 500 bp of upstream sequence was amplified from AR0382 genomic DNA using oTO1625-oTO1626. The *ADH1* terminator and *NEO* cassette were amplified from pTO169 using oTO1066-oTO668. Approximately 500 bp homologous to the region immediately downstream of *ALS4112* was amplified from AR0382 genomic DNA using oTO1627-oTO1628. The repair cassette was amplified from pTO337 using oTO18-oTO19 and transformed into *als4112* mutant strains to generate the complement strains.

To make *iff0675* mutant strain, the gRNA cassette was constructed with then Splice-On-Extension reaction by incorporating a 20-bp specific gRNA targeting the *IFF0675* gene in primers oTO841-oTO842. The repair template was amplified from plasmid pTO207 using oTO18-oTO19 and transformed into the Chicago4 strain to generate TO609.

### Genomic DNA Isolation

Yeast cells were cultured overnight in liquid YPD at 30°C, then collected by centrifugation and resuspended in a lysis buffer comprising 2% Triton X-100, 1% SDS, 100 mM NaCl, 10 mM Tris-Cl, and 1 mM EDTA. The cells were disrupted using bead-beating and the released DNA was first extracted with phenol-chloroform isoamyl alcohol (25:24:1) and then with chloroform extraction. Subsequently, the DNA was treated with RNase A followed by an additional precipitation step. It was finally resuspended in water for further use as PCR templates or sequencing.

### *C. auris* Transformation

The transformation of *C. auris* was performed using a transient Cas9 expression method as previously described ^41^. Briefly, transformation repair cassettes were amplified from the assembled plasmids. The Cas9 expression cassette was amplified from pTO135 using primers oTO143 and oTO41. For sgRNA targeting specific loci, expression cassettes were amplified from pTO136, utilizing overlap-extension PCR to modify the gRNA sequence. All linear PCR products were purified with the Zymo DNA Clean & Concentrator kit (Zymo Research, cat no. D4034,) following the manufacturer’s instructions.

To prepare electro-competent *C. auris* cells, cells were cultured in liquid YPD at 30 °C overnight with gentle shaking. Cells were then collected by centrifugation, resuspended in TE buffer containing 100 mM lithium acetate, and incubated for 1 hour at 30 °C with continuous agitation. DTT was added to the cells with a final concentration of 25 mM, followed by an additional 30-minute incubation at 30 °C. After centrifugation at 4 °C, the cells were washed once with ice-cold water and then with ice-cold 1 M sorbitol. The cells were finally resuspended in ice-cold 1 M sorbitol and either kept on ice for immediate use or aliquoted and stored at −80 °C for future transformations.

To perform electroporation, 45 μL of competent cells were mixed with 500-1000 ng of each PCR-amplified cassette, Cas9, sgRNA, and the repair template, and then added to a pre-chilled 2 mm gap electroporation cuvette. Cells were then electroporated with a Bio-Rad MicroPulser Electroporator, following the pre-defined PIC protocol (2.0 kV, 1 pulse). After electroporation, cells were recovered in 1 M sorbitol, resuspended in 10 ml YPD, and allowed to outgrow at 30 °C with continuous rotation for 2 hours. The outgrown cells were spread on selective media. Cells transformed with repair cassettes with the *NAT* marker or the *NEO* marker were selected on YPD with 200 μg/mL nourseothricin or 1 mg/mL G418, respectively and incubated at 30 °C for 2-3 days. Transformant colonies were verified for correct cassette integration by colony PCR using Phire Plant Direct PCR Master Mix (Thermo Fisher Scientific; F160) as per the manufacturer’s guidelines. Mutants were confirmed through at least three independent PCR reactions with unique primer sets targeting the mutation site and compared to the parental strains. For deletion mutants, gene-specific primers confirmed the absence of ORF amplification and proper integration of the repair cassette at the correct genomic locus.

### Keratinocyte adhesion assay

Human N/TERT keratinocytes ^35^ were cultured in keratinocyte-SFM medium (Gibco, ref. 10724-011, supplied with 30 μg/ml bovine pituitary extract, 0.2 ng/ml epidermal growth factor, and 0.31 mM CaCl_2_) to 100% confluence in 96-well plates and inoculated with 100 µl of diluted overnight *C. auris* culture (OD_600_=0.03). At the same time, 100 uL of diluted *C. auris* culture was inoculated into an empty 96-well plate, spun down at 1000 rpm for one minute, and images were taken to determine the input cell counts. For the adhesion assay, *C. auris* cells in the keratinocyte wells were spun at 1000 rpm for one minute and the plates were incubated at 37 °C with 5% CO_2_ for 1 hour to allow for adhesion. After incubation, non-adherent yeast was removed by washing three times with PBS. The cells were then fixed with 4% formaldehyde and stained with a FITC-conjugated anti-Candida antibody (Meridian Bioscience, cat. no. B65411F, 1:1000) to label *C. auris* cells, while Hoechst (Thermo Fisher Scientific, ref. 62249, 1:1000) and CellMask Deep Red (Thermo Fisher Scientific, cat. no. H32721, 1:5000) were used to stain keratinocyte nuclei and cell bodies, respectively. Plates were imaged using a BioTek Lionheart FX microscope at 10X magnification and adherent *C. auris* cells on keratinocytes were quantified using a custom CellProfiler ^68^ imaging pipeline developed by our lab. To count input cells, images were pre-processed in Fiji ImageJ software (version 2.14.0) ^69^ using a custom macro that segmented cells in the brightfield images using edge detection and generated a binary mask from the segmented cells. The pre-processed images were then quantified using CellProfiler to generate cell counts for the input. The percentage of adherent cells relative to input cells was used to assess the ability of each *C. auris* strain to adhere to keratinocytes.

To pretreat *C. auris* cells with serum for keratinocyte adhesion assay, 200 µL of Chicago4 cells cultured overnight at 30 °C were harvested and resuspended in 450 µL PBS with 50 µL of either anti-NDV-3A serum or anti-alum control serum. The cell suspensions were incubated on a rotator for 30 minutes at room temperature. After incubation, the cells were washed three times with PBS and diluted into keratinocyte culture medium for the keratinocyte adhesion assay.

### *C. auris* internalization assay

Keratinocytes were seeded in a CELLSTAR® Tissue Culture Plates (Greiner Bio-One, cat. no. 655090) and cultured until 100% confluent. *C. auris* Chicago4 cells were diluted to an OD_600_ of 0.03 in keratinocyte culture medium and applied to keratinocytes for 6 h at 37 °C. The cells were then washed three times with PBS and fixed with 4% formaldehyde for 10 minutes, followed by three washes with PBS. Cells were then blocked with 5% bovine serum albumin in PBS for 20 min. After blocking, cells were incubated with the FITC-conjugated anti-Candida antibody (Meridian Life Science, cat. no. B65411F, 1:1000) for 2 hours to label extracellular *C. auris* cells, followed by three washes with PBS. Keratinocytes were then permeabilized with 0.1% TritonX-100 for 30 minutes and washed three times with PBS. The total *C. auris* cells were then stained with 10% calcofluor white (Sigma-Aldrich, cat. no. 18909) for 10 min, followed by three washes with PBS. The images were captured on a BioTek Lionheart FX automated microscope.

### NF-**κ**B activation

Heat-killed *S. aureus* USA300 cells were prepared by incubating at 65°C for 30 minutes. *C. auris* Chicago4 cells were diluted to an OD_600_ of 0.03 in keratinocyte culture medium. To test NF-κB activation, keratinocytes were treated with heat-killed *S. aureus* cells, depleted zymosan (InvivoGen, cat. code tlrl-zyd, 100 ug/ml), or diluted *C. auris* cells at 37 °C for 6 h. After incubation, the culture medium was removed. The cells were then washed three times with PBS and fixed with 4% formaldehyde for 10 minutes, followed by another three washes with PBS. Keratinocytes were then permeabilized with 0.1% TritonX-100 for 30 minutes, washed three times with PBS, and blocked with 3% BSA plus 5% goat serum (Invitrogen, cat. no. 10000C) in PBS for 30 minutes. After blocking, keratinocytes were incubated with a Rabbit anti-NF-κB p65 antibody (Cell Signaling Technology, cat. no. 8242S, 1:800) diluted in the blocking solution for 2 hours, followed by three washes with PBS. Alexa Fluor 594-conjugated goat anti-rabbit secondary antibody (Thermo Fisher Scientific, A-11037, 1:500) and PierceTM Hoechst 33342 solution (ThemoFisher, 1:1000) diluted in the blocking buffer was added for 1 hour to mark NF-κB p65 and nuclei. After staining, cells were washed three times with PBS. The images were captured on a BioTek Lionheart FX automated microscope. Images from four different fields were taken for each condition to quantify NF-κB nuclear translocation.

To quantify NF-κB nuclear activation, we measured the integrated intensity of NF-κB signals within the nucleus using CellProfiler. The mean integrated intensity of nuclei under mock treatment was used as the baseline. Nuclei with integrated intensities above this baseline were classified as NF-κB-positive. The proportion of NF-κB-positive nuclei relative to the total nuclei was calculated to evaluate NF-κB activation for each condition. Quantification was performed on 5,037 cells for mock treatment, 4,674 cells for heat-killed MRSA treatment, 4,812 cells for deleted zymosan treatment, and 5,163 cells for *C. auris* treatment.

### *Agrobacterium tumefaciens*-Mediated Transformation (AtMT)

AtMT was performed as previously described ^41^. Briefly, *A. tumefaciens* strain pTO131 (EHA 105 containing pTO128) was cultured overnight in liquid LB media with kanamycin at 30 °C. The cells were collected by centrifugation, washed with sterile water, and then resuspended in liquid Induction Medium (IM) supplemented with 100 μM acetosyringone (3’,5’-dimethoxy-4-hydroxyacetophenone, AS) to a final OD_600_ of 0.15 and incubated at room temperature with constant shaking for 6 hours. Meanwhile, recipient *C. auris* Chicago4 (TO318) cells grown overnight at 30 °C in YPD were collected by centrifugation and resuspended in sterile water to an OD_600_ of 1.0. Equal volumes of prepared *A. tumefaciens* and *C. auris* cells were mixed and spread on solid IM Agar with AS, followed by incubation at 23 °C for 4 days. After incubation, cells were collected in liquid YPD and washed three times by centrifugation at 2000 rpm for 5 minutes to separate the fungal cells from the bacteria. Aliquots of the washed culture were then plated on YPD containing 200 μg/mL nourseothricin and 200 μg/mL cefotaxime, and incubated at 30 °C for 2 days. Transformant colonies were manually transferred to 96-well plates, grown in YPD overnight at 30 °C, then preserved by adding 50% glycerol to each well and storing the plates at −80 °C.

### High-throughput keratinocyte adhesion assay

The high-throughput assay was based on the above described keratinocyte adhesion assay with modifications. A total of 1,595 insertional mutants in the Chicago4 strain background were arrayed and individually assayed in 96-well plates. Each plate contained 92 individual wells of mutants and 4 wells of Chicago4 strain as controls. Mutant arrays were cultured from glycerol stocks on solid YPD agar at 30 °C. The resulting colonies were then used to inoculate 200 μL YPD cultures in 96-well plates and grown overnight at 30 °C. For the adhesion assay, *C. auris* cells from each well were transferred into 96-well plates containing keratinocytes at 100% confluence using a 96-well microplate replicator (RePad 96 long pin, REP-001, Singer Instruments, United Kingdom), with a single pinning. Input cells were prepared in the same way. The subsequent steps were carried out according to the keratinocyte adhesion assay.

For imaging, at least six different fields were captured. A ratio of adhesive cells was calculated by dividing the count of the attached cells remaining after washing by the count of the input cells for each captured field. A z-score was calculated for each mutant by subtracting the proportion of adhesive cells from the average proportion of adhesive cells amongst all the mutants and dividing by the standard deviation amongst all the mutants on that plate. Mutants with a z-score more negative than −1.8 were considered to have significantly reduced adhesion.

### AtMT Transgene Insertion Site Identification

Identification of transgene insertion sites was performed as described previously ^41^. Briefly, genomic DNA was isolated from the two mutants of interest, pooled, and sequenced by Illumina sequencing. Library preparation, quality control, and Whole Genome Sequencing were performed by SeqCenter (Pittsburgh, PA, USA). Libraries were prepared using the Illumina Nextera kit, and sequencing was performed on the NextSeq platform to produce 150 bp paired-end reads. The sequencing data was analyzed on the Galaxy web platform public server (usegalaxy.org) ^70^. Read quality was evaluated with FastQC, and reads were trimmed using Trimmomatic ^71^ with a Phred score cutoff of 20. The processed reads were aligned to a linearized reference of pTO128 (pPZP-Nat) using the Burrows-Wheeler Aligner (BWA-MEM) ^72^ with parameters set for a minimum seed length of 50 and a bandwidth of 2. Soft-clipped sequences corresponding to the genomic DNA neighboring the T-DNA integration sites, were extracted from the aligned BAM file using the extractSoftClipped script from SE-MEI (https://github.com/dpryan79/SE-MEI). These sequences were then mapped to the *C. auris* B8441 reference genome (NCBI GCA_002759435.2) using BWA-MEM with default settings to identify integration locations. Each T-DNA integration site in the sequenced mutants was verified by Sanger sequencing.

### Adhesin protein structure

Protein sequences for Als4112 and Iff0675 from different clades were obtained from FungiDB ^73^. The Iff0675 protein sequence from the Chicago4 strain was identified by sequencing the PCR product amplified with primers (oTO2170/oTO2171) targeting the *IFF0675* genomic locus, along with its upstream and downstream regions. The signal peptide domain was identified using automatic annotations from the UniProt database ^74^. The number of tandem repeats was manually counted, and repeat logos were generated using WebLogo ^75^. The GPI anchor region was determined using NetGPI 1.1 ^76^.

### RNA Extraction

RNA extraction was performed using a formamide extraction method ^77^. Briefly, cells cultured overnight at 30°C were harvested by centrifugation and frozen at −80 °C before processing. To extract RNA, cell pellets were thawed at room temperature and resuspended in 100 μL of FE Buffer, containing 98% formamide and 0.01 M EDTA. The we added 50 μL of 500 μm RNase-free glass beads to the cell suspension and the mixture was homogenized for 30 seconds three times using a BioSpec Mini-Beadbeater-16 (Biospec Products Inc., Bartlesville, OK, USA). The resulting cell lysate was then clarified by centrifugation to remove any cell debris, and the supernatant was collected as the crude RNA extract. This crude extract was purified with the Qiagen RNeasy Mini Kit (Qiagen, ref. 74104), following the manufacturer’s instructions. Samples were then treated with Invitrogen RNase-free DNase (Qiagen, cat no. 79254) to remove genomic DNA. The integrity and purity of the RNA were verified using a Nanodrop and by agarose gel electrophoresis before proceeding to downstream applications.

### RT-qPCR

RNA was isolated from *C. auris* cells cultured overnight in YPD at 30 °C and used to synthesize cDNA using the iScript cDNA synthesis kit (Bio-Rad Laboratories, cat no. 1708890) following the manufacturer’s instructions. The generated cDNA served as the template for qPCR reactions, which were set up using the Power-Up SYBR Green Master Mix (Thermo Fisher Scientific, cat. A25741) in accordance with the manufacturer’s protocol. Amplification was carried out using a Bio-Rad CFX Opus 384 Real-Time PCR System. Primers specific to the target genes were designed using NCBI Primer-BLAST ^78^ with the *C. auris* B8441 genome assembly (NCBI GCA_002759435.2) as a reference. *ACT1* was amplified with primers oTO359 and oTO360 and *ALS4112* was amplified with primers oTO448 and oTO449.

### ECM slide adhesion assay

An overnight culture of the RFP-tagged AR0383 strain was diluted to an OD_600_ of 0.05 in YPD medium. The ECM-coated slide from the ECM Select^®^ Array Kit Ultra-36 (Advanced BioMatrix, cat. no. 5170) was washed once with 5 mL PBS, followed by YPD medium, before use. The slide was then incubated with 5 mL of the diluted *C. auris* cells in the provided slide tray for 2 hours at room temperature with constant agitation. After incubation, the slide was washed three times with PBS and fixed with 4% PFA for 10 minutes at room temperature. The fixed slide was washed three times with PBS, and images of different ECM spots were captured using the Texas Red channel on the BioTek Lionheart FX microscope. The number of cells attached to each ECM spot was counted with CellProfiler.

### In situ adhesion to ECM of murine skin tissue

Uninfected frozen skin tissue specimens were cut to 8 μm using a cryostat and used for subsequent *in situ* adhesion assay. Each specimen was incubated with diluted overnight *C. auris* chicago4 cells (OD600=0.03) for 1.5 hours at room temperature and then washed three times with PBS to remove unattached cells. The specimens were fixed with 4% PFA for 10 minutes, followed by three washes with PBS. They were then stained with antibodies against laminin (Millipore Sigma, cat. no. L9393, 1:200), collagen I (Thermo Fisher Scientific, cat. no. MA1-26771, 1:200), or collagen III (Fisher Scientific, cat. no. PIMA542628, 1:100) and *C. auris* cells (Meridian Life Science, cat. no. B65411F, 1:1000). Images were taken with the BioTek Lionheart FX microscope.

### Adhesion competition assay on fibronectin- or laminin-coated coverslips

Overnight cultures of the RFP-tagged AR0383 strain and the *als4112* mutant strain in the AR0383 background were diluted to an OD*600* of 0.05 in YPD medium and mixed in a 1:1 ratio. 1 mL of the mixed culture was added to 24-well plates containing fibronectin- (Thermo Fisher Scientific, cat. no. NC0705540) or laminin-coated (Thermo Fisher Scientific, cat. no. NC0597572) coverslips. The cells were briefly centrifuged at 1000 rpm for 1 minute to promote contact with the surface, then incubated for 1 hour at room temperature to allow for adhesion. Before washing, the actual input ratio of the two strains was determined by imaging the coverslips in both the brightfield and Texas Red channels. After the 1-hour incubation, the coverslips were washed three times with PBS to remove non-adherent cells and fixed with 4% PFA for 10 minutes at room temperature. The fixed coverslips were then washed three times with PBS. Post-wash images were captured using the brightfield and Texas Red channels to determine the ratio of adherent cells. The ratio of adherent cells was normalized to the input ratio, and the normalized adherence was calculated, setting the adhesion of the *als4112* mutant strain to one.

### Epicutaneous Murine colonization

The murine epicutaneous infection was performed as previously described with minor modifications ^28^. Infection was performed using C57BL/6J mice at 7 weeks of age (Jackson Laboratories), which were in the telogen phase of hair growth. Mouse dorsal hairs were shaved and depilated one day before infection. For infection, *C. auris* strains were cultured overnight at 30 °C, washed once with PBS, and resuspended at a concentration of 1×10^9^ cells/ml in PBS using a hemocytometer. 1×10^8^ cells of *C. auris* were placed on a patch of sterile gauze and attached to the shaved skin of individual mice with a transparent occlusive plastic dressing (Tegaderm; 3M). The actual inoculum of each *C. auris* culture used for infection was determined by colony-forming units (CFUs) on YPD + ampicillin + gentamycin plates. Mice were sacrificed after being exposed to *C. auris* at the indicated time points. For CFU analysis, dorsal skin tissue underneath the gauze was harvested, weighed, and digested with 0.25 mg/ml liberase in 500 ul PBS for 1 hour and 45 minutes at 37 °C and 5% CO_2_. Digested dorsal skin tissue was homogenized with a bead beater for 30 seconds x 4 times. The supernatant of each homogenized sample was plated on YPD + ampicillin + gentamycin plates to determine the CFUs recovered from each mouse. The number of CFU was determined after 48 h of incubation at 30 °C. The actual fungal burden for each mouse was determined by normalizing the CFUs recovered from each mouse by the weight of the recovered dorsal skin tissue and actual delivered inoculum.

For immunofluorescence, the harvested skin tissues were fixed for 1 hr in cold 3.7% paraformaldehyde, washed with PBS, then incubated overnight in 30% sucrose at 4°C, before embedding in optimum cutting temperature compound (VWR, cat. no. 25608-930). 8 μm frozen tissue specimens were cut using a cryostat for subsequent staining. Specimens were first blocked with 20% normal goat serum in PBS for 20 minutes at room temperature. Primary antibodies diluted in 5% normal goat serum in PBS were applied and incubated overnight at room temperature. *C. auris* cells were detected with the FITC-conjugated anti-*Candida* antibody (Meridian Bioscience, cat. no. B65411F, 1:500), epidermis was marked by keratin 14 staining (BioLegend, cat. no. 906004, 1:1000). Specimens were then washed three times in PBS and incubated with secondary antibodies (Invitrogen, cat. no. A-21437) diluted in 5% normal goat serum in PBS (1:300) for 1 hour at room temperature in the dark. After removing the secondary antibody, Hoechst staining solution (Thermo Fisher Scientific, ref. 62249, 1:1000) was applied for 5 minutes to label nuclei. Samples were washed three times in PBS, then mounted with Vectashield for imaging. Images were taken using a Zeiss Observer D1 Inverted Fluorescence Microscope. In some cases, fluorescent images were processed using the Auto-Blend feature in Adobe Photoshop CS6 to automatically maximize image sharpness across multiple focal planes. H&E staining was performed by the University of Michigan Orthopaedic Research Laboratories Histology Core and images were taken with the BioTek Lionheart FX microscope.

### *Ex vivo* Human Skin colonization

Human skin colonization was performed as previously described ^28^. Briefly, samples were collected from patients who underwent reconstructive surgeries through an IRB-exempt protocol, and incubated as described ^19^, with paraffin wax as a barrier between skin surface and media. 10^5^ *C. auris* cells were then applied to the skin surface and incubated for 24 hours before processing for CFU analysis. Rinsed samples were compared to unrinsed samples to determine fungal adhesion.

### Intravenous Murine Infection: Inoculum Preparation, Mice Infection, and Histopathology

One day before infection, *C. auris* strains were cultured in 10 mL of YPD broth and incubated overnight in a shaker at 30 °C and 200 rpm. The next day, overnight yeast cultures were washed three times with 1X Dulbecco’s Phosphate Buffered Saline (DPBS) by centrifugation at 4000 rpm for 10 min at 4°C, followed by final resuspension in 10 mL of 1X DPBS. Yeast cells were counted using a hemocytometer, and the cell densities for each strain were adjusted to 2.5 × 10L cells/mL (5 × 10L cells/0.2 mL) using 1X DPBS.

Outbred ICR CD-1 mice (n = 12 per strain), aged 4–6 weeks, were immunosuppressed via intraperitoneal (i.p.) injection of 200 mg/kg cyclophosphamide and subcutaneous injection of 250 mg/kg cortisone acetate two days prior to infection. To prevent bacterial superinfection, enrofloxacin was added to the drinking water at a concentration of 50 μg/mL starting on the day of immunosuppression and continued for two weeks.

*C. auris* strains were injected via the tail vein, and the survival of infected mice was monitored for 21 days. Individual mouse weights were recorded on days 0, 4, and 7 post-infections to evaluate weight loss as a marker of disease progression. On day 7 post-infection, two mice from each group were euthanized, and their kidneys and hearts were harvested for histopathological examination. The harvested organs were fixed in 10% zinc-buffered formalin, embedded in paraffin, and sectioned. Tissue sections were stained with H&E at Harbor-UCLA Medical Center and imaged using an Olympus microscope.

### Growth Curve

Overnight cultures of *C. auris* were diluted to an OD_600_ of 0.01 in YPD medium. A total of 200 μL of the diluted culture was added to each well of a 96-well plate, with YPD medium alone used as a control. The plate was incubated with continuous shaking at 500 rpm at 30 °C. Growth was monitored using the BioTek LogPhase 600 plate reader, which measured OD600 every 10 minutes for 48 hours.

### Adhesion to BSA- or collagen-coated plastic surfaces

96-well plates were coated with 50 μL of collagen I (Sigma-Aldrich, cat. no. CC300, 100 μg/mL) in 10 μM HCl/PBS or collagen III (Advanced Biomatrix, cat. no. 5021, 100 μg/mL) in 10 μM HCl/PBS or 5% BSA. Plates were incubated overnight at 4 °C and washed once with 100 μL PBS before use. For the adhesion assay, overnight *C. auris* cultures were diluted to an OD_600_ of 0.03 in YPD medium. 100 μL of the diluted cells was added to either collagen-coated or uncoated wells. The plates were centrifuged at 1000 rpm for 1 minute to settle the cells, then incubated for 1 hour at room temperature to allow adhesion. After incubation, unattached cells were removed by washing the wells three times with 100 μL PBS. A vacuum aspirator with an 8-channel manifold (BrandTech QuickSip aspirator) was used to aspirate the media between each wash. Each well was imaged in brightfield using the BioTek Lionheart FX automated microscope both before and after washing. Six predefined fields per well were captured to ensure that the same regions were imaged before and after washing. Images were processed in the same manner as the input cell images for the keratinocyte adhesion assay. The number of cells in each field was quantified using CellProfiler software. The percent of adhesive cells for each field was calculated by dividing the count of cells remaining after washing by the count of input cells.

### Biofilm assay on collagen-coated 96-well plate

96-well plates were coated as described above. For the biofilm assay, we followed the protocol from Uppaluri et al with modifications ^79^. Briefly, overnight *C. auris* cultures were diluted to an OD_600_ of 0.03 in RPMI (Corning, ref. 10-040-CV) with 2% glucose. A total of 200 μL of the diluted cells was added to collagen-coated or uncoated wells. Plates were centrifuged at 1000 rpm for 1 minute to settle the cells, then incubated at 37 °C for 1 hour to allow adhesion. After incubation, unattached cells were removed by washing the wells three times with 100 μL PBS. To develop biofilm, 200 μL of fresh RPMI + 2% glucose was added to each well, and the plates were incubated at 37 °C with constant shaking at 250 rpm for 24 hours. Following incubation, the medium was aspirated, and the wells were washed twice with 200 μL PBS, ensuring the biofilm remained undisturbed. Biofilm was then quantified with crystal violet stain. After the PBS washes, the wells were air-dried and stained with 60 μL of 0.4% crystal violet solution (Sigma Aldrich, C-3886) for 45 minutes. Excess stain was removed by washing the wells four times with 200 μL PBS. To extract the crystal violet, 120 μL of 95% ethanol was added to each well and incubated for 45 minutes. Then, 100 μL of the ethanol extract was transferred to a fresh 96-well plate to take OD_595_. For biofilm imaging, after the two PBS washes, the biofilms were fixed with 100 μL of 4% PFA for 10 minutes at room temperature, followed by three PBS washes. The wells were then stained with 100 μL of 10% calcofluor white stain (Sigma Aldrich, 18909) for 10 minutes at room temperature and washed three times with PBS. Biofilms were imaged using the DAPI channel of the BioTek Lionheart FX microscope.

### Adhesion to PE catheter

PE catheters (Fisher Scientific, 14-170-11E) were cut into 2.5 mm pieces and coated overnight in collagen I or III solution in a 24-well plate. The coated catheters were then washed once with PBS before use. Overnight *C. auris* cultures were diluted to an OD_600_ of 0.1 in YPD medium. Five pieces of either uncoated or collagen I- or -III-coated catheters were placed into 1 mL of the diluted *C. auris* suspension in 1.5 mL microcentrifuge tubes and rotated on a mixer for 1 hour to allow cell attachment to the catheters. After incubation, the catheter pieces were transferred to a 48-well plate and washed three times with PBS. The catheters were then moved to microcentrifuge tubes, and 250 µL of PBS was added to each tube. The tubes were vortexed for 3 minutes to release the attached cells into solution. Next, 200 µL of the PBS solution was transferred to a 96-well plate and centrifuged at 1000 rpm for 1 minute to pellet the cells. Both input and attached cells were imaged in brightfield using the BioTek Lionheart FX automated microscope, and cell counts were quantified using CellProfiler software. The percentage of cells attached to the catheters was calculated for quantification.

### Rat Catheter Colonization

The *in vivo* biofilm assay was conducted using a rat external jugular venous catheter model, as described previously ^80^. Briefly, an inoculum of 10^6^ cells/mL for each strain was introduced onto an internal jugular catheter, which was placed in 16-week-old, 400 g specific-pathogen-free Sprague–Dawley rats. The biofilm was allowed to develop over a 24-hour period. After 24 hours, the rats were sacrificed, and the catheter was removed for CFU analysis. 1 mL of sterile 0.85% NaCl was gently flushed through the catheter to remove blood and nonadherent *C. auris* cells. The distal 2 cm of the catheter was then excised and placed into a 1.5-mL tube containing 1 mL of sterile 0.85% NaCl. The tube was sonicated in a FS 14 water bath sonicator with a 40-kHz transducer (Fisher Scientific) for 10 minutes, followed by vigorous vortexing for 30 seconds. Serial 1:10 dilutions of the catheter fluid were prepared and plated onto Sabouraud dextrose agar. Plates were incubated at 35 °C for 24 hours to get viable fungal colony counts.

### Statistics and Reproducibility

Graphical representations of numerical data were created using GraphPad Prism (v.10.4.1) for macOS. Statistical analyses of data were also conducted with GraphPad Prism. Unless specified otherwise, all experiments were conducted with a minimum of three independent biological replicates, and data are shown as means ± standard deviation of the means, with each data point representing an individual biological replicate. Microscopy and photography images are representative of at least two experiments that showed similar results. Statistical significance was determined using Student’s t-test or one-way ANOVA followed by Dunnett’s test for multiple comparisons. Correlation was determined using Pearson correlation. Survival analysis differences were analyzed using the Mantel-Cox log-rank test. \**P* ≤ 0.05; \*\**P* ≤ 0.01; \*\*\**P* ≤ 0.001; \*\*\*\**P* ≤ 0.0001, ns: *P* > 0.05.

### Ethics Statement

Mice were housed in the University of Michigan or The Lundquist Institute pathogen-free animal facility until infection, and all protocols were approved by and in compliance with the guidelines established by the IACUC at the respective institutions.

## Supplemental Figure Legends

**Fig. S1.** Quantification of NF-LB nuclear translocation under each condition was determined by the proportion of NF-κB-positive cells, identified by nuclei with integrated intensities above the mock treatment baseline. Values represent the mean ± s. d., calculated from cell images captured across four technical replicate wells. Statistical differences was determined using one-way ANOVA with Dunnett’s multiple comparisons test; \*\*\*\**P* ≤ 0.0001; ns: *P* > 0.05.

**Fig. S2. (A)** The growth of the *C. auris* wild-type from different clades, Δ*als4112*, and *ALS4112* complement strains at 30 °C in liquid YPD medium was assessed by measuring OD_600_ over a 48-hour period. **(B)** The transcript levels of *ALS4112* and *IFF4109* in Chicago4, Δ*als4112*, and complement strains was measured by qRT-PCR. The presence or absence of *IFF4109* and *IFF4110* in the genomes of Chicago4, Δ*als4112*, and complement strains was determined by colony PCR. Values represent the mean ± s. d., calculated from three biological replicates. Statistical differences were assessed using one-way ANOVA with Dunnett’s multiple comparisons test; \*\*\*\**P* ≤ 0.0001; ns: *P* > 0.05.

**Fig. S3. (A)** Adhesion of AR0382, AR1101, AR0383, and Chicago4 cells to uncoated or BSA-coated 96-well plates, measured by the proportion of cells remaining attached after washing. **(B)** Biofilm formation of AR0382, AR1101, and Chicago4 on uncoated or specific collagen-coated 96-well plates was quantified with crystal violet staining. Images of biofilm stained with calcofluor white were shown. Scale bar, 50 μm. Values represent the mean ± s. d., calculated from three biological replicates. Statistical differences were determined using unpaired t test (A) or one-way ANOVA with Dunnett’s multiple comparisons test (B); \*\**P* ≤ 0.01; \*\*\**P* ≤ 0.001; \*\*\*\**P* ≤ 0.0001; ns: *P* > 0.05.

**Table S1.** Strains used in this study.

| Strain Name | Genotype | Source |
| --- | --- | --- |
| TO33 | AR0382 | <sup>67</sup> |
| TO38 | AR0387 | <sup>67</sup> |
| TO39 | AR0388 | <sup>67</sup> |
| TO41 | AR0390 | <sup>67</sup> |
| TO703 | Clade I | This Study |
| TO704 | Clade I | This Study |
| TO32 | AR0381 | <sup>67</sup> |
| TO326 | AR1099 | <sup>67</sup> |
| TO326 | AR1100 | <sup>67</sup> |
| TO327 | AR1101 | <sup>67</sup> |
| TO34 | AR0383 | <sup>67</sup> |
| TO35 | AR0384 | <sup>67</sup> |
| TO328 | AR1102 | <sup>67</sup> |
| TO329 | AR1103 | <sup>67</sup> |
| TO707 | Clade III | This Study |
| TO709 | Clade III | This Study |
| TO712 | Clade III | This Study |
| TO36 | AR0385 | <sup>67</sup> |
| TO37 | AR0386 | <sup>67</sup> |
| TO52 | AR0931 | <sup>67</sup> |
| TO330 | AR1104 | <sup>67</sup> |
| TO331 | AR1105 | <sup>67</sup> |
| TO315 | Chicago1 | 5 |
| TO316 | Chicago2 | 5 |
| TO317 | Chicago3 | 5 |
| TO318 | Chicago4 | 5 |
| TO705 | Clade IV | This Study |
| TO708 | Clade IV | This Study |
| TO710 | Clade IV | This Study |
| TO711 | Clade IV | This Study |
| TO713 | Clade IV | This Study |
| TO53 | AR1097 | 67 |
| At pTO131 | *Agrobacterium tumefaciens* EHA105 harboring pTO128 (pPZP-NAT) | 41 |
| TO670 | Chicago4 *tnALS4112* (B9J08_004112) | This Study |
| TO671 | Chicago4 *tnMUB1* (B9J08_003899) | This Study |
| TO672 | Chicago4 P1C8 | This Study |
| TO186 | AR0382 $\Delta als2582$(B9J08_002582)::NAT | 28 |
| TO187 | AR0382 $\Delta als4498$(B9J08_004498)::NAT | 28 |
| TO226 | AR0382 $\Delta als4112$(B9J08_004112)::NAT | 28 |
| TO233 | AR0382 $\Delta iff4100$(B9J08_004100)::NAT | 28 |
| TO234 | AR0382 $\Delta iff4109$(B9J08_004109)::NAT | 28 |
| TO235 | AR0382 $\Delta iff4098$(B9J08_004098)::NAT | 28 |
| TO236 | AR0382 $\Delta iff4110$(B9J08_004110)::NEO | 28 |
| TO247 | AR0382 $\Delta iff1531$(B9J08_001531)::NAT | 28 |
| TO248 | AR0382 $\Delta iff4892$(B9J08_004892)::NAT | 28 |
| TO249 | AR0382 $\Delta iff1155$(B9J08_001155)::NAT | 28 |
| TO250 | AR0382 $\Delta iff4451$(B9J08_004451)::NAT | 28 |
| TO251 | AR0382 Δiff0675(B9J08_000675)::NAT | 28 |
| TO261 | AR0382 Δscf1(B9J08_001458)::NAT | 28 |
| TO568 | AR0382 Δals4112(B9J08_004112)::NAT<br>+ NAT::ALS4112<sub>AR0382</sub> (B9J08_004112)<br>NEO | This Study |
| TO554 | AR1101Δals4112(CJI96_0005138)::NAT | This Study |
| TO565 | AR1101 Δals4112(CJI96_0005138)::NAT<br>+ NAT::ALS4112<sub>AR0382</sub> (B9J08_004112)<br>NEO | This Study |
| TO444 | AR0383 Δals4112(CJI97_004175)::NAT | This Study |
| TO567 | AR0382 Δals4112(CJI97_004175)::NAT<br>+ NAT::ALS4112<sub>AR0382</sub> (B9J08_004112)<br>NEO | This Study |
| TO445 | Chicago4 Δals4112(CJJ07_003308)::NAT | This Study |
| TO666 | AR0382 Δals4112(CJJ07_003308)::NAT<br>+ NAT::ALS4112<sub>AR0382</sub> (B9J08_004112)<br>NEO | This Study |
| TO609 | Chicago4 Δiff0675(CJJ09_002636)::NAT | This Study |
| TO436 | AR0382 pTEF1-ALS4112 NAT | 28 |
| TO134 | AR0383 ENO1::ENO1-RFP+NAT | 41 |

**Table S2.** Plasmids used in this study.

| Name | Description | Source |
| --- | --- | --- |
| pTO135 | pENO1-CaCas9-tCyc1, AMP | 41 |
| pTO136 | pADH1-tRNA-Ala-gRNA-tracrRNA-HDV-tAgTEF2, AMP | 41 |
| pTO139 | pUC19, AMP | 41 |
| pTO166 | als4112(B9J08_004112)::NAT, AMP | 28 |
| pTO169 | ACE2 NEO, AMP | 41 |
| pTO337 | *ALS4112<sub>AR0382</sub> (B9J08_004112) NEO, AMP* | This study |
| pTO207 | *Iff0675(CJJ09_002636)::NAT, AMP* | 28 |

**Table S3.**
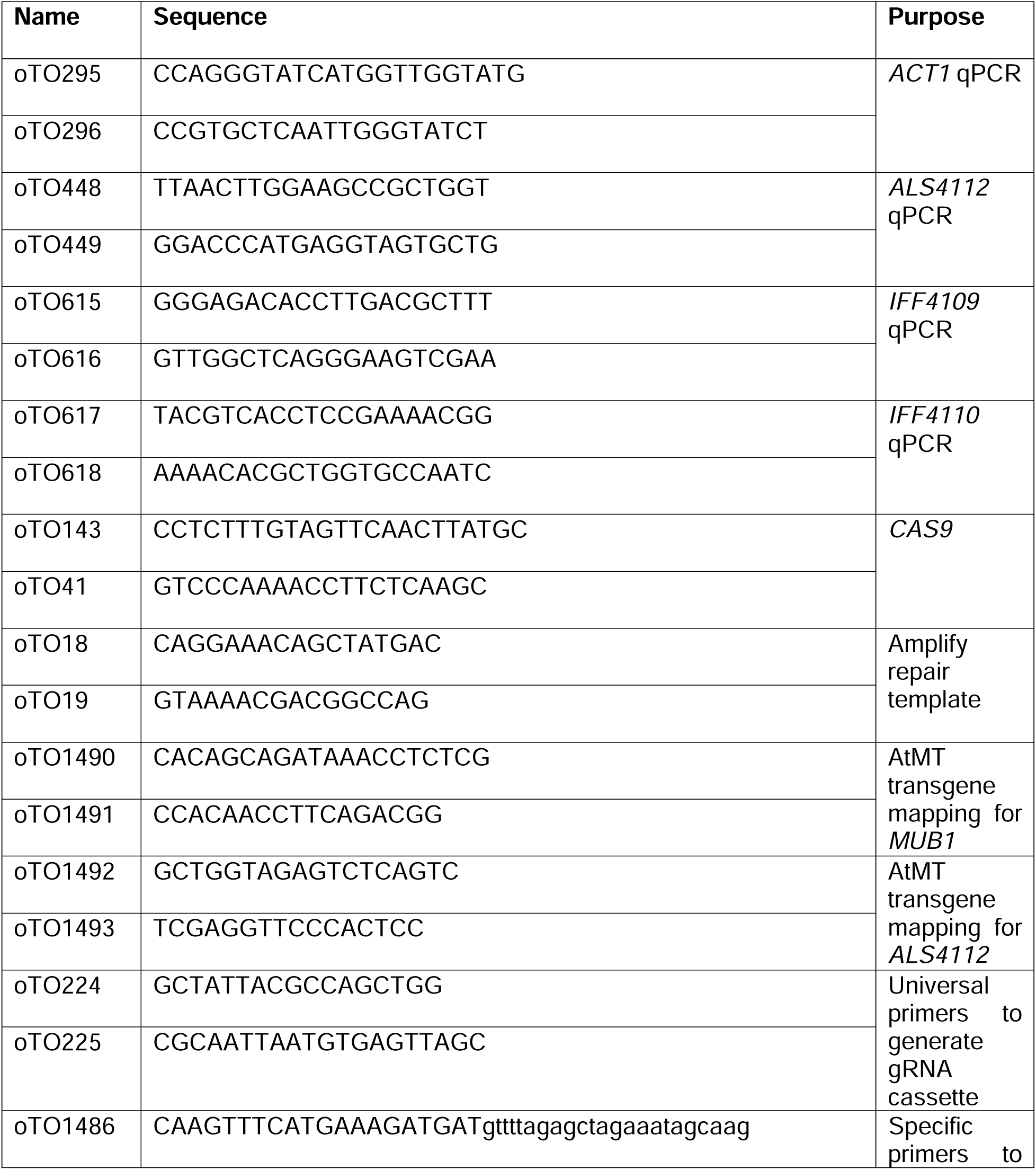
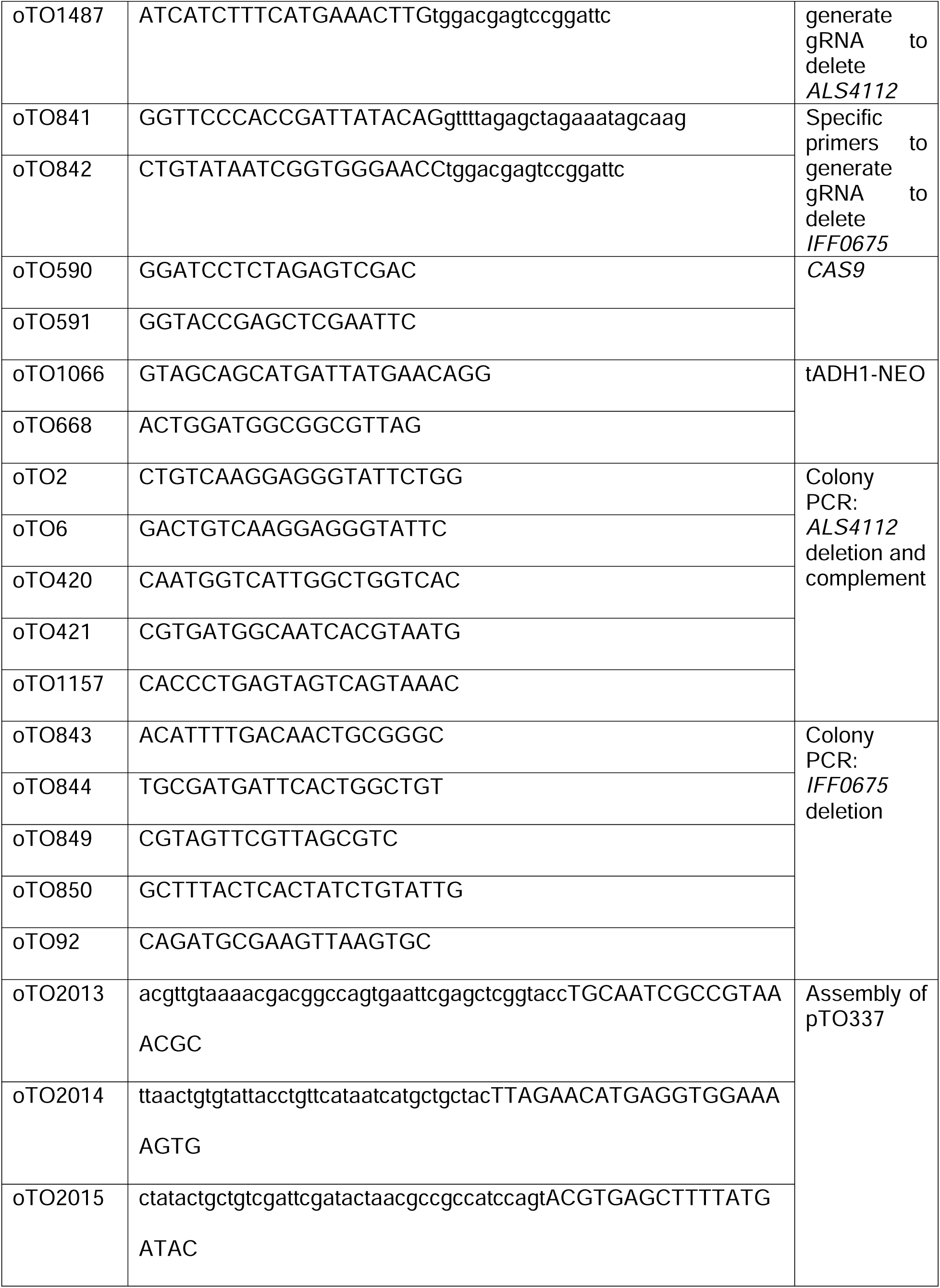
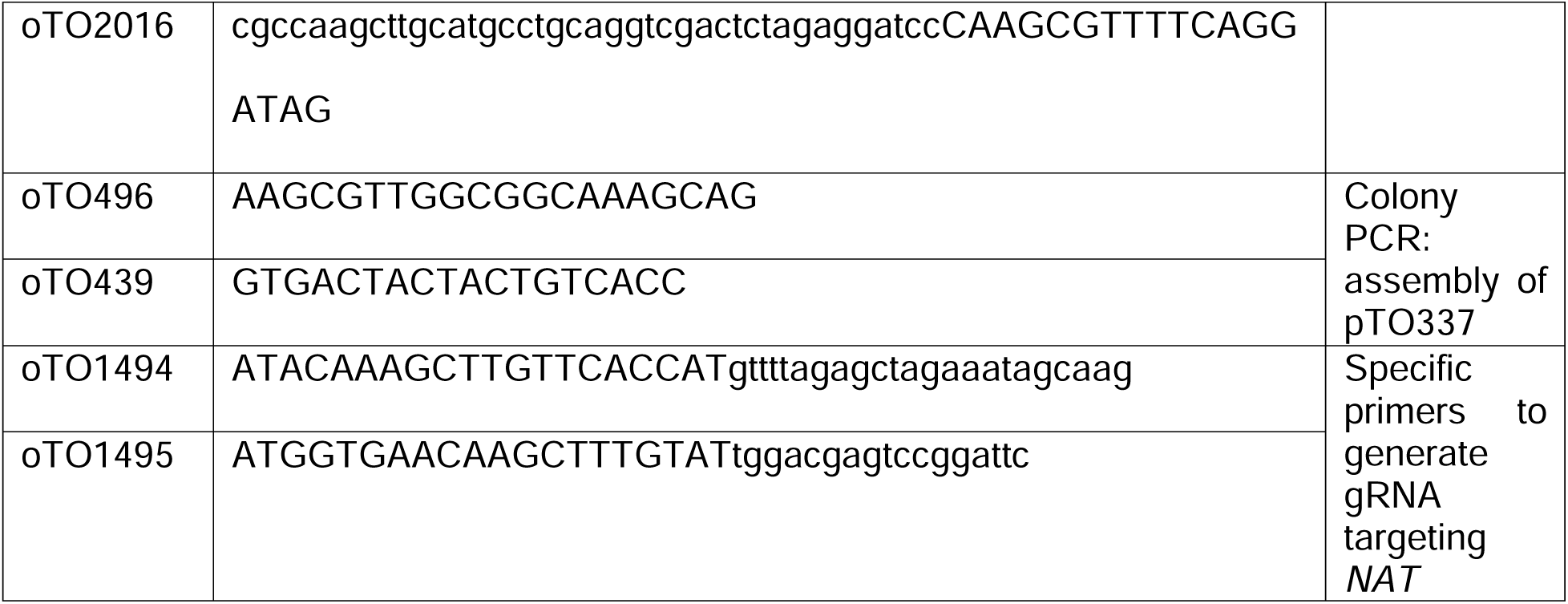
Oligos used in this study.

## Acknowledgments

We thank Michael J. McFadden for consultation on image analysis, Carey Dombecki and the UM infection prevention team and Mary K. Hayden for sharing *C. auris* strains. We are grateful for histological support from Emma Snyder-White (through the Michigan Integrative Musculoskeletal Health Center Core).

This work was supported by the National Institutes of Health grant T32AI007413, F32AI181164, and Michigan Pioneer Fellows Program Fund to G.Z.; NIH grant R21AI169186, U19AI181767, and Burroughs Wellcome Fund Investigator in the Pathogenesis of Infectious Diseases Award to T.R.O.; NIH grant F31AI169823 to D.J.S.; NIH grant R01AI145939 to J.E.N.; R01AR080654 and R01AR065409 to S.Y.W.; NIH grant K24AR076975 to J.M.K.; NIH 1K01AI180591, NIH NCATS UCLA CTSI KL2TR001882, and American Heart Association Award #938451 to S.S.. Research reported in this publication was supported by the National Institute of Arthritis and Musculoskeletal and Skin Diseases of the National Institutes of Health under Award Number P30 AR069620.

The content is solely the responsibility of the authors and does not necessarily represent the official views of the National Institutes of Health.

## Author contributions

Conceptualization: G.Z., T.R.O.; Funding acquisition: G.Z., T.R.O., D.J.S., J.E.N., S.Y.W., J.M.K., S.S.; Investigation: G.Z., J.L., N.A.V., J.A.E.A., B.X., R.Z., E.M., C.J.J., D.Q., H.H., K.D., L.A.H., S.S., D.J.S.; Methodology: G.Z., J.L., N.A.V., J.A.E.A., B.X., R.Z., E.M., C.J.J., D.Q., H.H., K.D., L.A.H., S.S., D.J.S.; Project administration: G.Z., T.R.O.; Supervision: T.R.O., D.A., J.E.N., S.S., A.S.I., M.L.K., A.C.A., S.Y.W., J.M.K., Visualization: G.Z., T.R.O.; Writing – original draft: G.Z., T.R.O.; Writing – review and editing: G.Z., J.L., J.A.E.A., R.Z., C.J.J., H.H., N.D.V., A.S.I., D.A., J.E.N., S.S., M.L.K., S.Y.W., T.R.O.

## Data and materials availability

constructs generated in this study will be provided for research purposes upon request. All remaining data are available in the main text or the supplementary materials.

