## Supplementary figures and images for "*Candida auris* skin colonization is mediated by Als4112 and interactions with host extracellular matrix proteins"

### s1

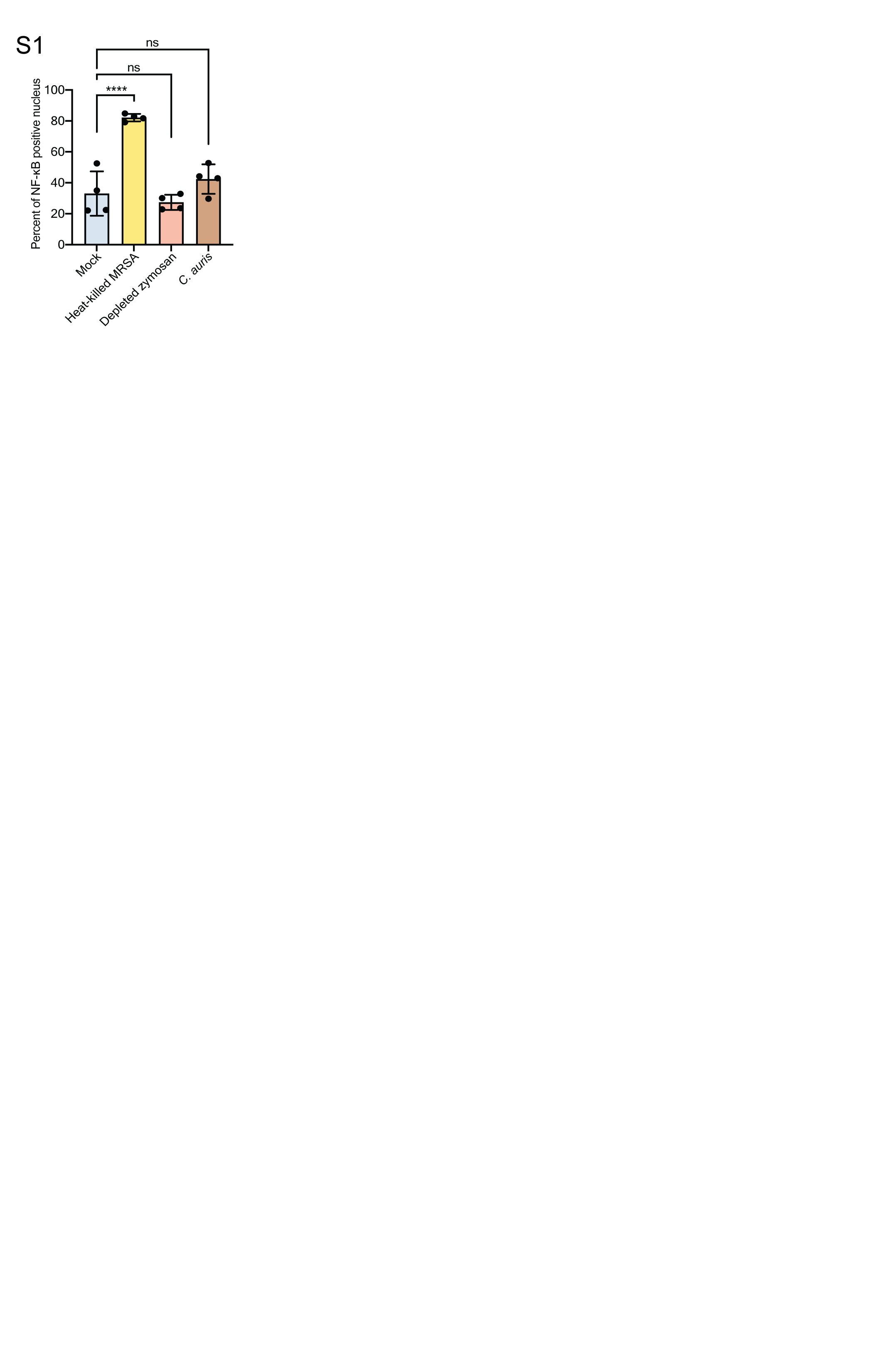

### s2

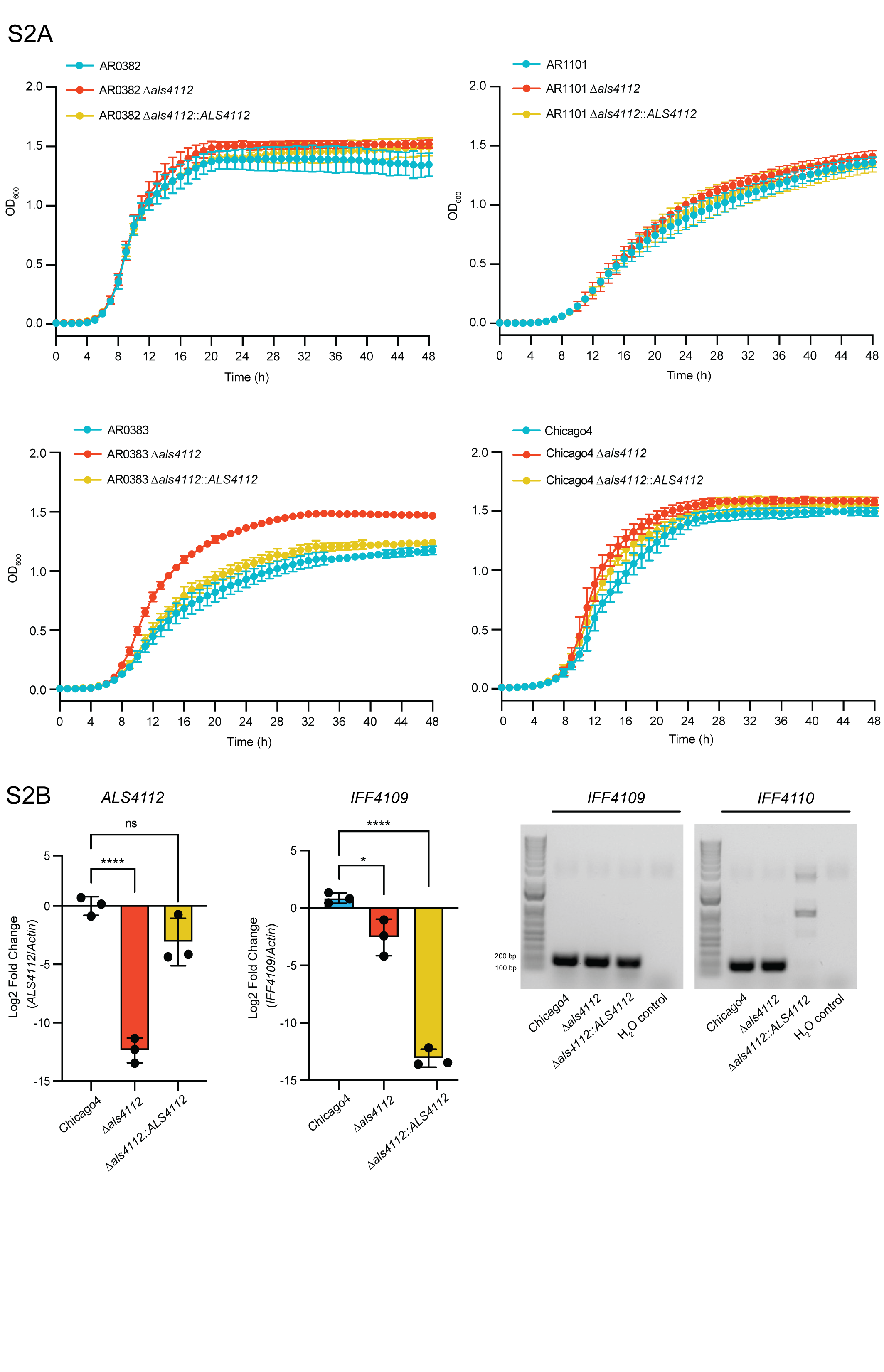

### s3

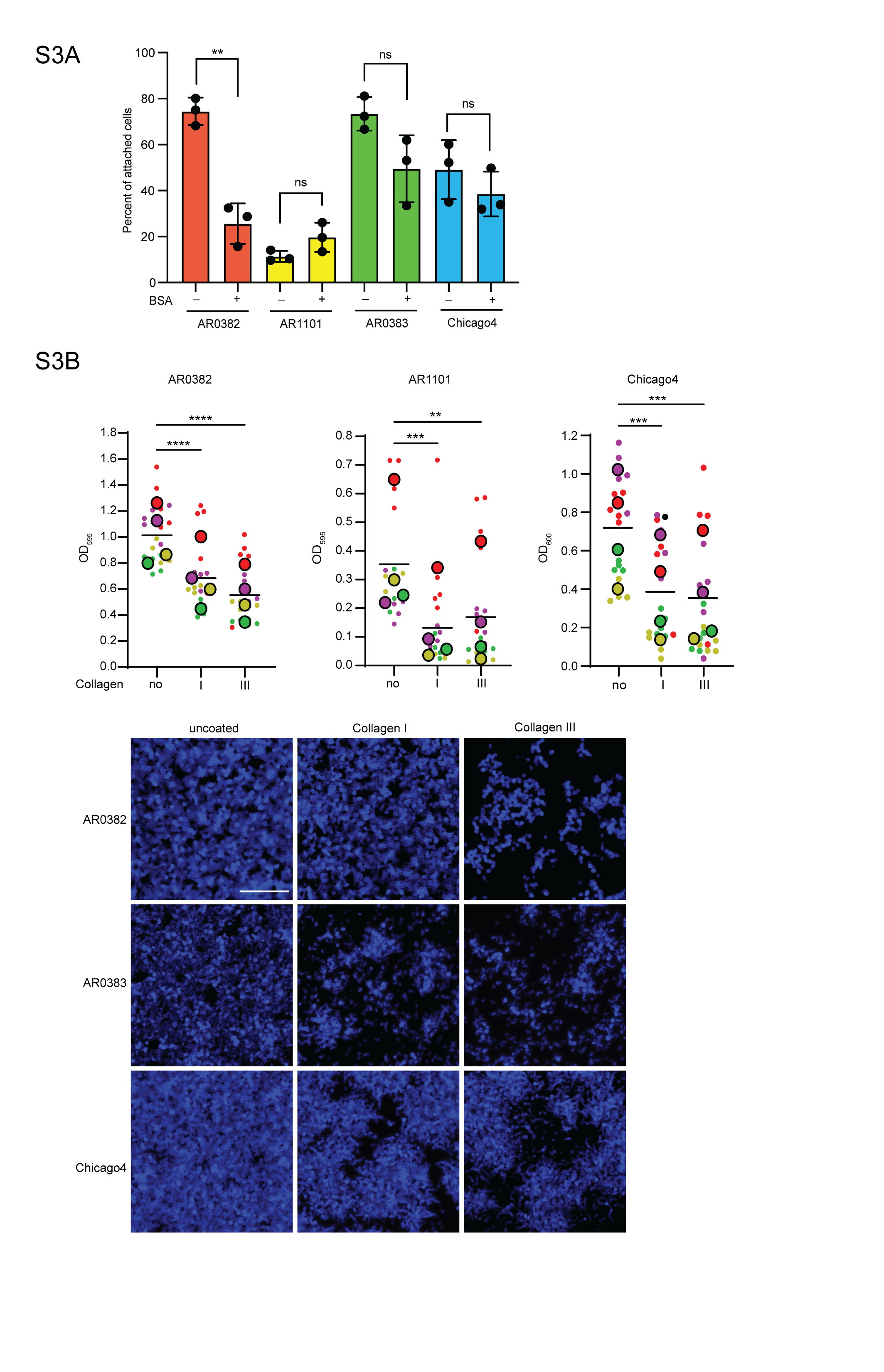
